# Donor tissue-resident memory-like T and NK cells generate a transient peripheral chimerism in lung transplant recipients, potentially protective from chronic lung allograft dysfunction

**DOI:** 10.1101/2023.06.16.543190

**Authors:** Ramon Bellmàs-Sanz, Anna-Maria Hitz, Bettina Wiegmann, Jenny F. Kühne, Evgeny Chichelnitskiy, Theodore S. Kapellos, Kim-Alina Bläsing, Jana Keil, Kerstin Beushausen, Wiebke Sommer, Kristian Händler, Matthias Becker, Kevin Baßler, Lisa-Marie Horn, Danny Jonigk, Mark Greer, Axel Haverich, Joachim L. Schultze, Fabio Ius, Gregor Warnecke, Christine S. Falk

## Abstract

Lung transplantation (LTx) is the only definite treatment option of patients suffering from end-stage lung disease. Long-term outcome is hampered by chronic allograft dysfunction resulting from poorly defined immune mechanisms. In this study of 97 lung recipients, we demonstrate dynamic changes of T, B and NK cell subsets early after lung transplantation with a selective decrease in memory CD4^+^ and CD8^+^ T cells accompanied by a relative increase in NK cells. Simultaneously, donor-derived T and NK cells were detected in recipient blood (n=44) immediately after LTx, persisting for three weeks. Donor T and NK cells displayed a CD69^+^ but CD103^-^CD49a^-^ CD25^-^ phenotype, which was shared by T and NK cells in lung perfusion solutions. In order to uncover the origin of these donor T and NK cells, the immune compartment of human lung explant tissue, i.e. trachea and parenchyma, was analyzed and it revealed three major subsets: classical circulating CD69^-^CD103^-^CD49a^-^ T and NK cells, CD69^+^CD103^+^CD49a^+^ tissue-resident memory (TRM) T and NK cells and CD69^+^CD103^-^CD49a^-^ TRM-like T and NK cells. Single-cell RNA sequencing confirmed the presence of TRM-like subsets with unique features, which reflected the phenotypes of donor T and NK cells and created a transient chimerism in recipient blood. Higher frequencies of donor T cells within the first three weeks showed a tendency for protection from chronic lung allograft dysfunction (CLAD) two years after transplantation although the correlation analyses did not reach statistical significance. To the best of our knowledge, we show for the first time that a transient chimerism is established within the first weeks after lung transplantation by donor TRM-like T and NK cells, which may contribute to protection from chronic lung allograft dysfunction (CLAD) development.

**Single Sentence Summary:** *Donor TRM-like T and NK cells cause a transient chimerism in lung recipients and potentially contribute to protection from CLAD*.

## Introduction

Lung transplantation (LTx) often remains the only definite treatment option for patients with end-stage lung diseases. Despite the advances in surgical techniques, organ preservation and immunosuppression, the survival rates for lung transplant recipients are still poor compared to other solid organ transplantations (1). Primary graft dysfunction (PGD) at an early stage and chronic lung allograft dysfunction (CLAD) at later stages are the major conditions limiting survival. PGD develops during the first 72 hours after transplantation and is characterized by edema and hypoxemia accompanied by immune cell infiltration in the transplanted lung (2). CLAD typically develops months to years after transplantation and manifests in two major forms: the bronchiolitis obliterans syndrome (BOS) and restrictive allograft syndrome (RAS) (3). Immune-dependent mechanisms such as alloreactive effector T cells, natural killer (NK) cells and donor-specific antibodies (DSA) are considered to play a major role in the initiation of both PGD and CLAD (4). Thus, monitoring of the longitudinal dynamics of T and NK cell subsets early following lung transplantation represents a feasible strategy to study their impact on PGD and CLAD and to identify potential biomarkers candidates.

The lung as an interface organ is permanently exposed to environmental particles and/or pathogens that can potentially induce inflammation, allergy or severe respiratory tract infections. To combat this permanent threat, the lungs are equipped with a complex set of immune cells that promote protection against airway pathogens and maintain tissue homeostasis. Alveolar macrophages, dendritic cells, and T cells represent the most abundant pulmonary immune cells, although NK cells and other innate lymphoid cells (ILC) play as well major roles in lung immunity (5). Numerous immune cell subsets (e.g. naïve/memory/effector CD4^+^ and CD8^+^ T cells, NK cells) have lately been identified in the human lung via use of single-cell RNA sequencing (6, 7).

The human lung harbors a large population of a specialized non-circulating T cell subset known as tissue-resident memory T cells (TRM) (8). TRM T cells confer optimal protection against previously encountered respiratory pathogens *in situ*, and several studies have shown that the lung is populated with TRM T cells specific for airway pathogens such as influenza (9). TRM T cells are CD45RO^+^CCR7^-^ T cells with a unique transcriptional profile that distinguish them from circulating effector memory (EM) T cells (10, 11). The key marker to discriminate TRM T cells from circulating counterparts is CD69, a type II C-lectin receptor shown to sequestrate and downregulate sphingosine-1-phosphate receptor 1 (S1PR1), thus avoiding tissue egress towards sphingosine-1-phosphate (S1P) gradients (12). CD103 (αE integrin) and CD49a (α1 integrin) are also hallmarks of tissue residency and might play an important role in anchoring TRM T cells onto the epithelium (13). In addition, TRM T cells have been characterized by the upregulation of inhibitory receptors (i.e. PD-1, CD101), by an altered chemokine receptor composition (i.e. upregulation of CXCR6 and downregulation of CX3CR1) as well as enhanced levels of cytotoxicity-related gene transcripts (i.e. *GZMA, GZMB, GNLY, PFN1, IFNG, TNF*) even in steady/not-activated states (10, 11, 13).

Likewise, tissue-resident NK cells have recently been described in the human lung and are also characterized by the expression of CD69, CD49a and CD103 (14, 15). This tissue-resident phenotype has been associated to CD56^bright^ NK cells, whereas the majority of lung CD56^dim^ NK cells were found to be CD69^-^, and consequently proposed to represent mostly circulation-derived cells (16).

During transplantation, the immune cell compartment populating the donor lungs is transferred with the graft into the recipient. It is well known that certain lymphocyte populations present in the graft migrate into the periphery of the recipient, accounting for a transient chimerism in the blood (17). Nevertheless, neither the phenotype nor the clinical impact of these passenger donor lymphocytes has been studied in detail. The fact that the lung is highly enriched with tissue-resident T and NK cells raises the question whether passenger donor lymphocytes exhibit tissue-resident properties or, otherwise, display a circulatory phenotype. Although passenger donor lymphocytes are potential candidates for the development of graft versus host disease (GVHD), they might be beneficial by contributing to tolerance, such as the killing of allo-specific T cells by donor NK cells (18). A recent publication suggested that the persistence of TRM T cells in the bronchioalveolar lavage (BAL) correlates with a better outcome of lung transplant recipients (19).

In this study, we aimed to investigate the characteristics and origin of donor-derived T and NK cells detected in the periphery of lung transplant recipients (n=97) and to determine their impact on clinical outcome. We show dynamic changes of T, B and NK cells early after lung transplantation with a selective decrease in memory CD4^+^ and CD8^+^ T cells accompanied by a relative increase in NK cells directly after transplantation. Simultaneously, donor-derived so-called passenger T and NK cells could be detected in the periphery of every recipient (n=44) immediately after transplantation. Of note, these donor T and NK cells expressed higher levels of CD69 compared to recipient cells but did lack CD49a and CD103 tissue residency, as well as activation markers. This peculiar phenotype was shared by T and NK cells in lung perfusion solutions at the end of preservation, indicating that these cells may represent a mobile immune compartment of the lung. This hypothesis could be confirmed in tissues of the lower respiratory tract, i.e. the trachea and lung parenchyma. Of note, the trachea contained almost exclusively CD69^+^CD103^+^CD49a^+^ tissue-resident memory T and NK cells referenced to as true-TRM. In contrast, lung parenchyma contained a mixture of these TRM T and NK cells, classical circulating CD69^-^CD103^-^CD49a^-^ T and NK cells and peculiar subsets of CD69^+^CD103^-^CD49a^-^ TRM-like T and NK cells. Since this characteristic TRM-like T and NK cell subset was also found in recipient blood as well as perfusion solutions, these cells may still be able to leave the lung – in contrast to true-TRM T and NK cells – and generate a transient chimerism in the recipient’s blood. Of note, upon *in vitro* activation donor T cells were superior in production of cytotoxic molecules, i.e. IFN-γ, and their higher frequencies within the first three weeks after LTx showed a tendency to be associated with freedom from CLAD two years after transplantation. Using single-cell RNA sequencing, we could confirm the existence of distinct TRM and TRM-like T cell subsets in lung parenchyma with specific expression profiles, which assign higher retention marker expression to TRM and higher cytotoxicity and higher chemokine expression to the TRM-like T cell subsets. To the best of our knowledge, we show for the first time that a transient chimerism is established in the first weeks following lung transplantation by donor TRM-like T and NK cells, which appears to be protective from the subsequent development of CLAD.

## Materials and methods

### Study design

97 patients that underwent lung transplantation at Hannover Medical School between 2011 and 2019 were included in this study. From each patient blood was collected at different time points: before transplantation (pre), directly after transplantation (D0), 24 hours after transplantation (D1) and three weeks after transplantation (D21). In addition, perfusion solution (perfusate) from 111 transplanted lungs was obtained. Lung parenchyma was collected from explanted lungs of transplant recipients, whereas trachea was obtained from donor lungs used for transplantation. Clinical and demographical characteristics of study participants are summarized in **Table 1**. The study was approved by the Hannover Medical School Ethics committee (n° 2500-2014 and n° 7414). All patients provided written informed consent before participation in the study.

**Table 1.** Patient demographics and clinical characteristics.

| Characteristics | Study cohort<br>(n=97) | HLA cohort<br>(n=44) | Perfusate<br>(n=111) |
| --- | --- | --- | --- |
| **Age at Tx (years)** | 48.3±13.9 | 48.1±13.5 |  |
| **Gender** |  |  |  |
| male | 46 (47.4%) | 21 (47.7%) |  |
| female | 51 (52.6%) | 23 (52.3%) |  |
| **Double/single Tx** |  |  |  |
| Double LTx | 95 (97.9%) | 43 (97.7%) |  |
| Single LTx | 2 (2.1%) | 1 (2.3%) |  |
| **Transplant indication** |  |  |  |
| COPD | 25 (25.8%) | 13 (29.5%) |  |
| IPF | 33 (34.0%) | 15 (34.1%) |  |
| CF | 22 (22.7%) | 9 (20.5%) |  |
| PHT | 4 (4.1%) | 2 (4.5%) |  |
| ReTx | 4 (4.1%) | 1 (2.3%) |  |
| Others | 9 (9.3%) | 4 (9.1%) |  |
| **PGD (Grade 2-3)** |  |  |  |
| 24h post Tx | 20 (20.6%) | 13 (29.5%) |  |
| 48h post Tx | 25 (25.8%) | 15 (34.1%) |  |
| 72h post Tx | 16 (16.5%) | 7 (15.9%) |  |
| **CLAD 2 years post Tx (n=73/39)** | 13 (17.8%) | 7 (17.9%) |  |
| BOS | 9 (12.3%) | 4 (10.3%) |  |
| RAS | 4 (5.5%) | 3 (7.7%) |  |
| **Organ preservation** |  |  |  |
| SOC | 65 (67.7%) | 30 (68.2%) | 79 (71.2%) |
| ESLP | 32 (33.3%) | 14 (31.8%) | 32 (28.8%) |
| **CCT (min)** | 494.5±117.2 | 497.7±125.0 | 480.0±112.4 |
| **Donor age (years)** | 46.9±15.7 | 49.7±15.7 | 45.6±16.0 |
| **Donor gender** |  |  |  |
| male | 46 (47.4%) | 21 (47.7%) | 50 (45.0%) |
| female | 51 (52.6%) | 23 (52.3%) | 61 (55.0%) |

BOS: Bronchiolitis Obliterans Syndrome; CCT: Cross Clamp Time; CF: Cystic Fibrosis; CLAD: Chronic Lung Allograft Dysfunction; COPD: Chronic Obstructive Pulmonary Disease; ESLP: Ex Situ Lung Perfusion; IPF: Idiopathic Pulmonary Fibrosis; PGD: Primary Graft Dysfunction; PHT: Pulmonary Hypertension; RAS: Restrictive Allograft Syndrome; ReTx: Retransplantation; SOC: Standard Of Care; Tx: Transplantation Age and CCT data are mean values ± standard deviation (SD). HLA cohort was a subgroup of the study cohort. No significant ( $p<0.05$ ) differences could be detected between both cohorts (unpaired t-test, Mann-Whitney test).

### Isolation of cells from blood and perfusates

Peripheral blood mononuclear cells (PBMCs) were isolated from blood by gradient separation using Biocoll separating solution (Biochrom). The resulting PBMCs where either used immediately for flow cytometry experiments or cryopreserved for further analyses. Absolute cell counts were calculated from whole blood using BD Trucount™ Tubes (BD Biosciences), following manufacturer’s instructions. Cells present in perfusates were treated with BD Pharm Lyse™ lysing solution (BD Biosciences) to lyse erythrocytes and subsequently stained for flow cytometry.

### Isolation of cells from lung tissue

Lung parenchyma and trachea tissue samples were digested using BD Dissociation reagent TuDOR (BD Biosciences) according to manufacturer’s instructions. The resulting single cell suspensions were filtered through a 70µm nylon filter and treated with BD Pharm Lyse lysing solution (BD Biosciences) to eliminate erythrocytes. The obtained single cell suspensions were either used immediately for flow cytometry experiments or cryopreserved for further analyses. For scRNA experiments, cells were additionally filtrated over 30μm cell strainer to reduce the duplets.

### Stimulation of cells

PBMCs were stimulated with phorbol 12-myristate-13-acetate (PMA) (50ng/ml) and ionomycin (2,5µg/ml) in tumor medium (RPMI 1640 medium supplemented with 2 mM L-glutamine, 1 mM sodium pyruvate, 100 U/ml Penicillin-Streptomycin and 10% FCS) for 15 hours. Brefeldin A (5µg/ml) was added into the wells 2 hours after the stimulation was initiated.

### Flow cytometry

All steps were performed as recommended by the guidelines for the use of flow cytometry in immunological studies (26). Samples were incubated at 4°C for 30 min unless otherwise indicated; all antibodies used for flow cytometry analyses are listed in **Table S3**. To assess cell viability cells were first stained for 15 min at room temperature with LIVE/DEAD Fixable Yellow Dead Cell Stain Kit (Thermo Fisher Scientific) in PBS. Cells were washed twice with PBS and incubated with antibodies for surface staining in FACS Buffer (0,1% NaN3, 2,5% FCS in PBS). Cells were washed once more prior to acquisition. For intracellular IFN-γ staining, cells were subsequently fixed and permeabilized with Intracellular Fixation/Permeabilization Buffer Set (eBioscience), washed once and incubated with IFN-γ antibody in Permeabilization buffer (eBioscience). Cells were then washed once again before acquisition. For staining with unconjugated HLA-antibodies, cells were first incubated with unconjugated donor-specific HLA antibodies and after one washing step they were stained with secondary fluorochrome-labelled goat anti mouse antibodies. Cells were acquired on a LSRII flow cytometer (BD Biosciences) using FACS Diva software (v8.0, BD Biosciences). Data was analyzed using FACS Diva software.

### Single cell RNA sequencing and bioinformatics

The scRNA libraries were generated using Seq-Well protocol as described before (20), with 20,000 single cells of each analyzed tissue cells loaded per array. The libraries were sequenced using NextSeq500 system with coverage of approximately 100.000 reads per cell.

### Processing of scRNA-seq raw data and bioinformatic analysis

The generated fastq files were applied into a data pre-processing pipeline (version 0.31, available at https://github.com/Hoohm/dropSeqPipe) followed by processing the data with R package ‘Seurat’ (v.3.0.0, (27)) for downstream analyses. Data were log normalized; top 2000 most variable features were identified using the “vst” method of the FindVaribaleFeature function of Seurat; variable features were z-transformed (scaled) and the influence of total UMI counts per cell was regressed out; scaled variable features were used as input to PCA; harmony was applied to the PCA embedding by correcting for the influence of the factor “LuT”; 30 harmony-corrected dimensions were used as input to UMAP. For cluster identification, a resolution of 0.6 was used, while the neighborhood graph was build based on 30 harmony-corrected dimensions. Gene sets were enriched using the AddModuleScore function of Seurat and visualized using the Nebulosa package. Package versions: Seurat_4.0.4; harmony_0.1.0 and Nebulosa_1.4.0.

The default setting for number of neighbors (k=20) were used. Marker genes for identified cell types/clusters were calculated using a Wilcoxon rank sum test for differential gene expression implemented in Seurat. The significance-threshold for marker genes was set to an adjusted p-value smaller than 0.1 and the logarithmic fold change cutoff to at least 0.25. In addition, the detected marker genes should have been expressed in at least 10% of the cells within the respective cell types/clusters.

To support cluster annotation, we analyzed for the enrichment of published gene signatures i.e. mature-TRM (19): *TMSB4X, CD7, IL32, RAD51D, CCL5, ZNF683, KLRD1, PFN1, CCR5, ACTB, NDUFA5, CD27, CD52, NKG7, GZMA, ITGA1, CXCR6, RUNX3*; TRM-like (19): *MLLT3, CACNA1A, PDZD3, PIG, ARSK, HSPA4, SLC25A37, COL6A3, SUV39H2, TTPAL, FAM107A, CDH6, YIPF4, LCMT2, BTBD9, SOX11, RIC3, DOCK7, TNFSF13B*; CD4^+^ Tregs (21): *ID3, SELL, STAT1, MAL, IFI6, LGALS3, MAF, TNFRSF18, FOXP3, IL2RA, CTLA4, TIGIT, PMCH, TNFRSF4, IFIT3*; CD8^+^ EM/resting TRM (21): *TCF7, AQP3, LGALS3, KLRB1, AMICA1, ANXA2, CRIP1, ITGA1, HOPX, GZMK, GNLY, MYO1F, CCL5*.

### Statistical analyses

Statistical analyses of the data were performed with GraphPad Prism v7.0 software (GraphPad Software Inc.). D’Agostino-Pearson omnibus normality test was calculated to assess data distribution. Parametric tests were performed when data was normally distributed, otherwise non-parametric tests were used. When comparing different time points between individuals, paired tests were used. The statistical test used in each analysis is indicated in the Figure legends.

Results were considered significant if p<0.05. Qlucore Omics Explorer (version 3.6, Qlucore) was used to generate principal component analysis (PCA) plots and heatmaps. The software was used to perform two-group or multigroup comparisons that identify the immune cell subsets that are most highly significantly different between the groups analyzed. For each analysis, the q-value used as a cut-off is indicated in the Figure legends. In paired analysis, influence from different baseline levels of patients was removed by selecting the annotation “patient” as eliminated factor. Missing values were given an estimated value calculated using the kNN algorithm provided by the software.

## Results

### Differential lymphocyte depletion in recipient blood directly after lung transplantation

The longitudinal dynamics of lymphocyte subsets were studied in a cohort of 97 patients who underwent lung transplantation at Hannover Medical School. From each recipient, blood samples were collected before (pre), directly after (D0), 24 hours (D1) and 3 weeks (D21) post transplantation and the lymphocyte composition in these samples was analyzed in detail by flow cytometry (**Fig. 1**). The frequencies of 11 immune cell subsets varied significantly between these early time points and samples from D0 and D1 clustered separately compared to pre-transplant and D21 samples, indicating that the main alterations in lymphocyte subsets occur during this early period (**Fig. 1A, B, Fig. S1A**). The proportions of T cells decreased directly after transplantation, mainly due to a decline in the frequency of CD4^+^ T cells (**Fig. 1C-D**), which resulted in a significantly altered CD4^+^/CD8^+^ ratio (**Fig. 1E**), altering the initial prevalence of CD4^+^ T cells before transplantation (**Fig. 1F**). In parallel, NK cell proportions showed a peak at D0 but returned to pre-transplant levels already at D1, whereas B cells levels were increased at D0 and D1 (**Fig. 1D**). At D21, lymphocyte subsets showed different frequencies compared to all other time points, though not returning to pre-transplant levels. Importantly, absolute cell numbers of all lymphocyte subsets declined at D0 and D1, especially CD4^+^ T cell numbers (**Fig. S1A**), explaining the modified lymphocyte proportions detected early after transplantation. Lung preservation, i.e. cold static (standard of care, SOC) *vs*. normothermic oxygenated preservation (*ex vivo* lung perfusion, EVLP) did not impact on lymphocyte alterations early after LTx as no differences in lymphocyte frequencies were observed between both preservation protocols (**Fig. S1B**).

**Fig. 1.**
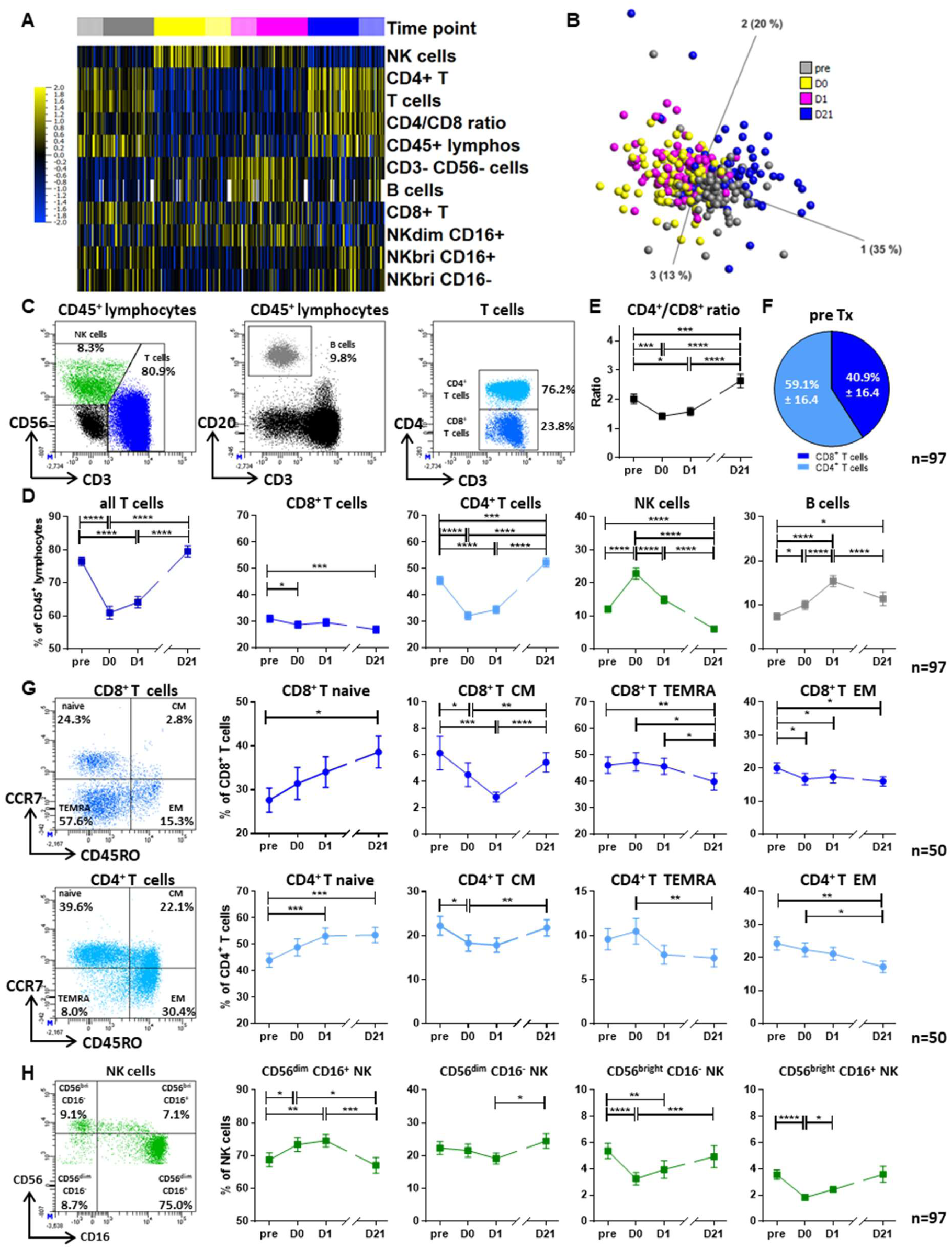
Early dynamic changes in lymphocyte subsets in blood after transplantation. The immune cells distribution in recipient blood was analyzed using flow cytometry before transplantation (pre), and directly (D0), 24 hours (D1) and 3 weeks (D21) after transplantation. Flow cytometry plots are representative of a pretransplant sample. **A**. Heatmap generated after performing a multigroup comparison between the analyzed time points (n=82) using a panel of 12 immune cell subsets characterized via flow cytometry. Patient was selected as an eliminated factor. With a q-value cut-off of 0.05, frequencies of 11 subsets were found significantly different among the groups. Subsets are ordered in the heatmap according to increasing q-values. The yellow to blue scale indicates the prevalence of each subset. Missing values are highlighted in green. **B.** Principle component analysis (PCA) performed including the 11 subsets whose frequencies differed significantly between time points. **C**. Representative flow cytometry plots describing the strategy used to identify CD8^+^ and CD4^+^ T cells, NK cells and B cells among CD45^+^ lymphocytes. **D.** Proportions of lymphocyte subtypes at the indicate time points (mean ± SEM, n=97, ANOVA calculated for n=82). **E**. Ratio between the proportions of CD4^+^ at CD8^+^ T cells at the indicated time points (mean ± SEM, n=97). **F**. Frequency of CD4^+^ and CD8^+^ T cells among T cells before transplantation (pre) (mean ± SD, n=97). **G.** Representative flow cytometry plots and proportions of CD8^+^ and CD4^+^ T cell subsets at the indicated time points (mean ± SEM, n=50, ANOVA calculated for n=38). Subsets were defined as naïve (CD45RO^-^CCR7^+^), CM (CD45RO^+^CCR7^+^), TEMRA (CD45RO^-^CCR7^-^) and EM (CD45RO^+^CCR7^-^). **H**. Representative flow cytometry plot and proportions of NK cell subsets at the indicated time points (mean ± SEM, n=97, ANOVA calculated for n=82). Subsets were defined as CD56^dim^CD16^+^, CD56^dim^CD16^-^, CD56^bright^CD16^-^ and CD56^bright^CD16^+^ cells. Statistical analysis: Repeated Measures ANOVA with Tukey multiple comparison test or Friedman test with Dunn’s multiple comparison test. Asterisks indicate p values (*p<0.05, **p<0.01, ***p<0.001, ****p<0.0001).

Next, the dynamics of naive and memory T cell and major NK cell subsets were determined in detail. CCR7^+^CD45RO^-^ naive T cell frequencies progressively increased after transplantation, accompanied by a decrease of CCR7^+^ CD45RO^+^ central memory (CM) and CCR7^-^CD45RO^+^ effector memory (EM) subsets in both CD8^+^ and CD4^+^ T cells (**Fig. 1G**). Recipient age had a minor effect in these changes (**Fig. S1C**). In NK cells, the proportion of CD56^dim^CD16^+^ NK cells increased directly after transplantation, whereas the CD56^bright^ subsets declined transiently (**Fig. 1H**). Taken together, LTx recipients show an immediate lymphopenic status accompanied by changes in the lymphocyte composition directly after lung transplantation, raising the question of whether these fluctuations may have an impact on clinical outcome and the balance between tolerance and rejection.

### Donor passenger T and NK cells display a memory phenotype and associations to clinical outcome

We hypothesized that the presence of donor passenger lymphocytes could be at least partially responsible for the substantial dynamics in lymphocyte proportions directly after LTx. The HLA mismatch between lung donors and recipients was utilized in 44 patients to quantify donor T and NK cells by staining with the respective donor HLA class I-specific antibodies by flow cytometry (**Fig. 2A**). Remarkably, donor T and NK cells were detected in the peripheral blood of all 44 patients immediately after LTx (**Table S2**), though with high individual variability. NK cells represented the major population of donor lymphocytes (**Fig. S2A**) and at D0, on average 16% of all circulating NK cells were donor-derived (**Fig. 2A**). In addition, donor T cells were also recovered at clearly detectable amounts with higher levels of donor CD8^+^ than CD4^+^ T cells (**Fig. 2A, B**). While donor T cells almost disappeared from recipient blood at D21, a significant proportion of donor NK cells was still detectable at levels above the detection limit of 1%. Interestingly, patients in the ESLP group showed a non-significant trend towards lower levels of donor T cells, implying that the preservation technique of the donor organ may have an impact rather on donor T cell than on NK cell frequencies (**Fig. S2B**). However, the presence of donor lymphocytes was independent of the cross-clamp time (CCT, **Fig. S2C**).

**Fig. 2.**
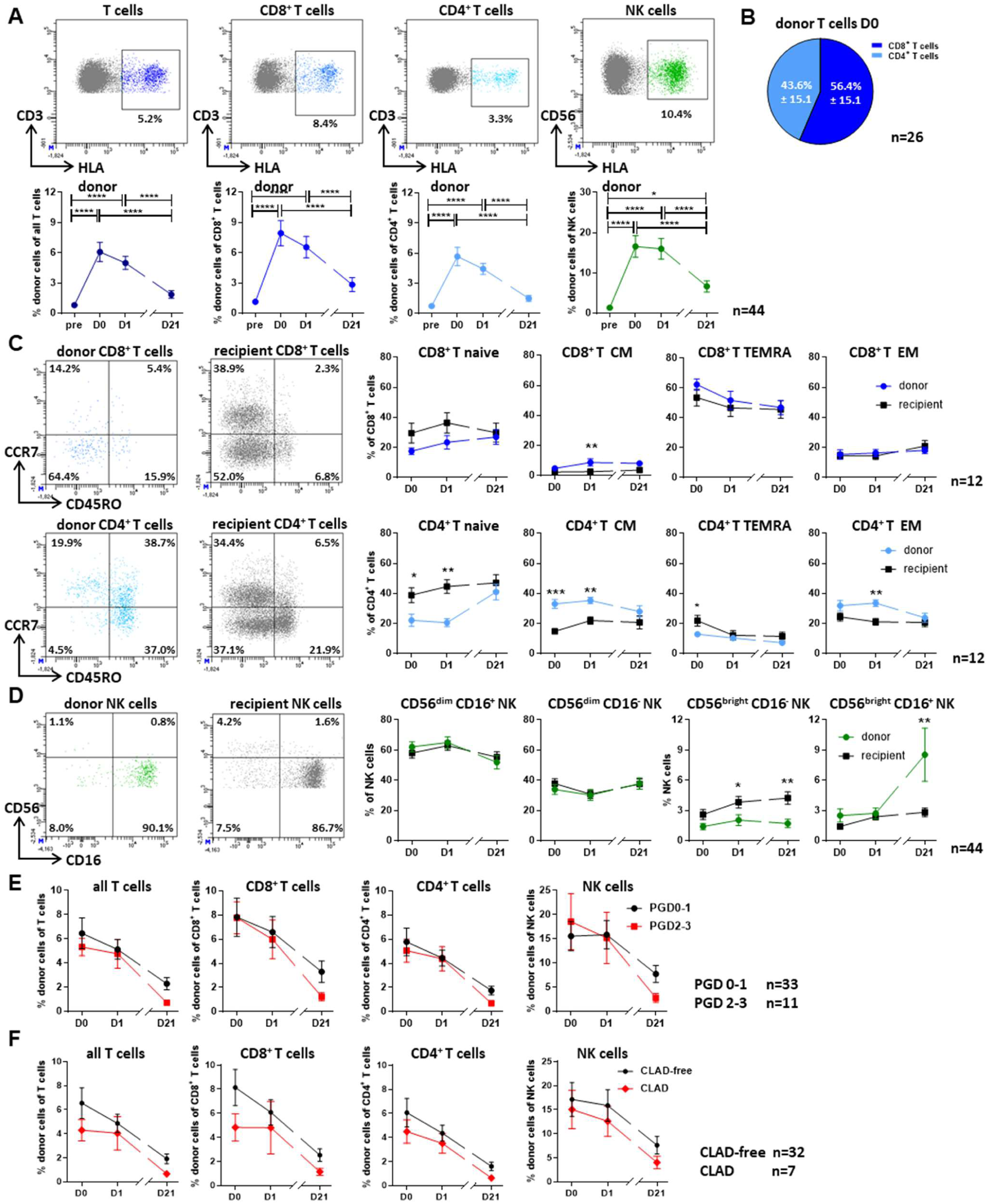
Specific subsets of donor T and NK cells are present in recipient blood after transplantation. **A**. Representative flow cytometry plots and frequencies of donor cells among all T cells, CD8^+^ T cells, CD4^+^ T cells and NK cells before transplantation (pre), and directly (D0), 24 hours (D1) and 3 weeks (D21) after transplantation (mean ± SEM, n=44). Donor cells were identified by staining donor HLA molecules in HLA-mismatched donor/recipient pairs. **B**. Frequency of CD4^+^ and CD8^+^ T cells among donor T cells directly after transplantation (D0) (mean ± SD, n=44). **C**. Proportions of CD8^+^ and CD4^+^ T cell subsets among donor and recipient T cells at the indicated time points (mean ± SEM, n=12). Subsets were defined as naïve (CD45RO^-^CCR7^+^), CM (CD45RO^+^CCR7^+^), TEMRA (CD45RO^-^ CCR7^-^) and EM (CD45RO^+^CCR7^-^). **D**. Proportions of NK cell subsets among donor and recipient NK cells at the indicated time points (mean ± SEM, n=44). Subsets were defined as CD56^dim^CD16^+^, CD56^dim^CD16^-^, CD56^bright^CD16^-^ and CD56^bright^CD16^+^ cells. **E**. Frequencies donor T cell subsets and NK cells in patients regarding their severity for primary graft dysfunction (PGD) at 24h after transplantation. Patients were divided between PGD0-1 (black, n=33) and PGD2-3 (red, n=11). Shown is mean ± SEM. **F**. Frequencies donor T cell subsets and NK cells in patients concerning their development of chronic lung allograft dysfunction (CLAD) at 2 years after transplantation. Patients were divided between CLAD-free (black, n=32) and CLAD (red, n=7). Shown is mean ± SEM. Statistical analysis: **A**. Friedman test with Dunn’s multiple comparison test. **C-F**. Two-way repeated measures ANOVA with Sidak multiple comparisons test. Shown are only the significant differences between study groups and not between time points. Asterisks indicate p values (*p<0.05, **p<0.01, ***p<0.001, ****p<0.0001).

Of note, donor-derived T and NK cells showed a different phenotype than their recipient counterparts. In the donor T cell pool, we detected higher proportions of central and effector memory T cells but lower proportions of naïve T cells compared to recipient T cells. This T cell subset distribution was especially pronounced for CD4^+^ T cells (**Fig. 2C, Fig. S2D**) and generally, independent of donor age (**Fig. S2F**). In turn, the vast majority of donor NK cells showed a CD56^dim^ phenotype, and the presence of CD56^bright^CD16^-^ donor NK cells was significantly reduced, compared to recipient NK cells (**Fig. 2D, Fig. S2E**). The increased presence of CD56^bright^CD16^+^ donor NK cells at D21 may be indicative of NK cell activation due to CD56 upregulation.

Next, we analyzed the potential impact of donor passenger T and NK cells on early and late clinical outcome after LTx. The presence of donor T and NK cells did not differ between patients without (grade 0-1) or with PGD grade 2-3 (**Fig. 2E**), supporting the established concept that PGD primarily results from ischemia/reperfusion injury. In contrast, there was a non-significant trend towards higher proportions of donor T cells, especially CD8^+^ T cells, in patients without CLAD, arguing for a beneficial effect of donor T cells on CLAD-free survival (**Fig. 2F**). This observation suggests that donor T cells in recipient blood may have a long-term protective role in LTx.

### Donor T and NK cells in recipient blood display a tissue-resident memory-like phenotype

The lung is populated by high numbers of tissue-resident memory (TRM) immune cells, which prompted us to investigate whether donor T and NK cells migrating out of the pulmonary graft into recipient blood may display TRM features. Thus, expression of the TRM markers CD69, CD103 and CD49a was analyzed in parallel in donor and recipient lymphocytes. Donor T and NK cells constantly displayed higher CD69 expression than recipient cells with significantly highest levels at D21 (**Fig. 3A**). However, the activation marker CD25 was barely expressed at D0 and D1 (**Fig. 3B**), arguing for CD69 as TRM rather than an activation marker. Later at D21, a small subset of donor T cells co-expressed CD25 (**Fig. 3B**), supporting our interpretation of CD69 early as TRM and only subsequently as an activation marker. Unexpectedly, donor T and NK cells were negative for the classical TRM markers CD103 and CD49a at all time points as were recipient T and NK cells (**Fig. 3C**). The lack of these TRM markers suggests that donor T and NK cells may not represent classical CD69^+^CD103^+^CD49a^+^ TRM T/ NK cells but rather CD69 single-positive TRM-like T cells, as defined by Szabo *et al*. (21).

**Fig. 3.**
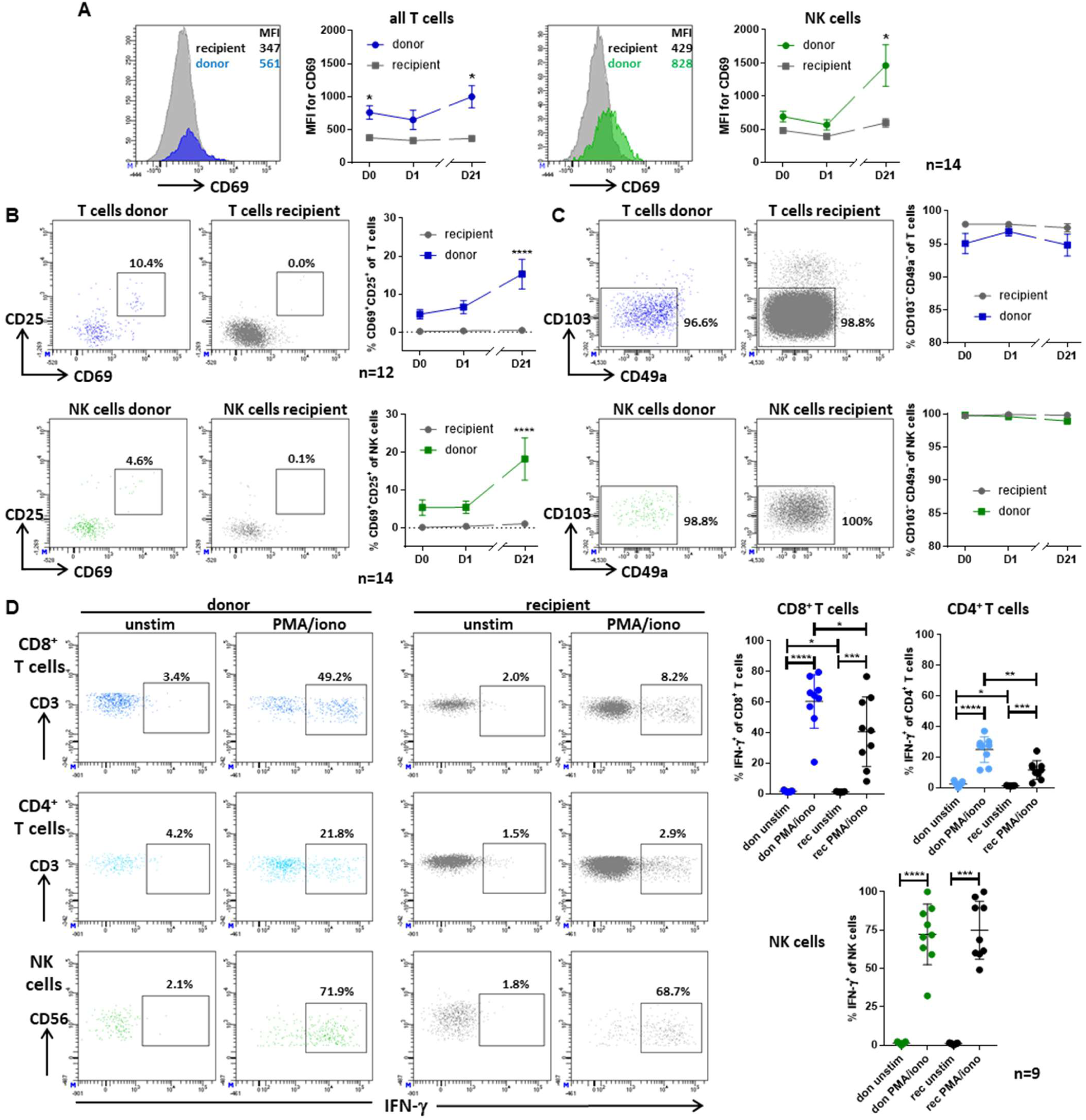
Donor T and NK cells express higher levels of CD69 than recipient cells but lack other markers of tissue residency. **A**. Representative histograms and graphs showing the median fluorescence intensity (MFI) for CD69 in donor and recipient T and NK cells directly (D0) 24 hours (D1) and 3 weeks (D21) after transplantation (mean ± SEM, n=14). **B**. Representative flow cytometry plots and frequencies of CD69^+^CD25^+^ T and NK cells among donor and recipient cells at the indicated time points (mean ± SEM, n=14). **C**. Representative flow cytometry plots and frequencies of CD103^-^CD49a^-^ T and NK cells among donor and recipient cells at the indicated time points (mean ± SEM, n=12). **D**. Representative flow cytometry plots and frequencies of IFN-γ^+^ cells among CD8^+^ T, CD4^+^ T and NK cells from donor and recipient origin obtained directly after transplantation (D0) that were cultured for 15h with PMA/ionomycin (PMA/iono) or left unstimulated (unstim) (mean ± SD, n=9). Statistical analysis: **A-C**. Two-way repeated measures ANOVA with Sidak multiple comparisons test. Shown are only the significant differences between study groups and not between time points. **D**. Paired t-test or Wilcoxon signed rank test between donor/recipient and PMA/iono/unstim pairs. Asterisks indicate p values (*p<0.05, **p<0.01, ***p<0.001, ****p<0.0001).

Next, the functionality of donor vs. recipient T and NK cells was compared at D1 after LTx in terms of cytokine (IFN-γ) induction. Donor CD8^+^ and CD4^+^ T cells produced higher amounts of intracellular IFN-γ than recipient T cells following non-specific stimulation with PMA/ionomycin (**Fig. 3D**). Moreover, the spontaneous IFN-γ release of T cells in culture was also higher in donor compared to recipient T cells. In contrast, no functional differences were observed between donor and recipient NK cells in that both were able to secrete high levels of IFN-γ upon stimulation.

Taken together, these results show that donor T and NK cells shared some but not all features with TRM cells and that donor T cells possess a higher functional capacity, i.e. IFN- γ secretion than their recipient counterparts.

### Lung perfusion solutions contain T and NK cells expressing CD69 but no other TRM- markers

During lung preservation, some immune cells are able to leave the lung into the perfusion solution, representing a source of pulmonary immune cells. Perfusate T and NK cells were immune phenotyped, focusing on lineage and TRM markers. T cells comprised the majority of perfusate lymphocytes although NK cells were also found at high frequencies with a mean of 31.7% (**Fig. 4A**). Unlike circulating blood T cells and similar to donor T cells, CD8^+^ T cells were present at higher frequencies than CD4^+^ T cells. Among CD8^+^ T cells, TEMRA T cells were the most frequent subset, while among CD4^+^ T cells, EM cells were the most prominent subpopulation (**Fig. 4A**). Of note, donor age had only a minor effect on the T cell subset distribution (**Fig. S3A**). In the NK cell compartment, CD56^dim^CD16^+^ cells comprised the majority of perfusate NK cells, whereas CD56^bright^ were present at very low frequencies (**Fig. 4A**).

**Fig. 4.**
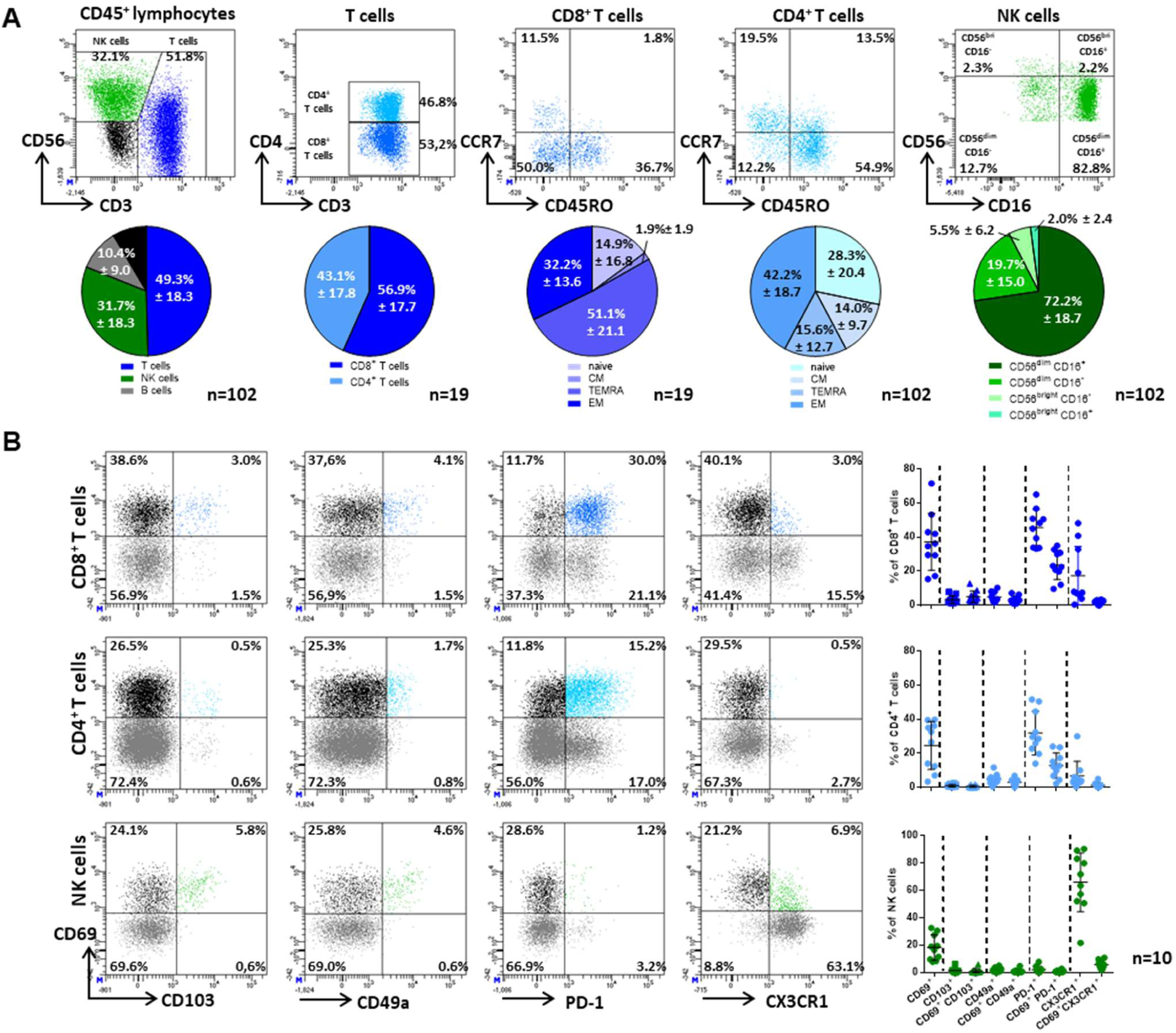
Perfusate cells express CD69 but lack other TRM markers. **A**. Representative flow cytometry plots and frequencies of used T cells, NK cells and B cells (n=102) and the subsets of T cells (n=19) and NK cells (n=102) in lung perfusates. T cell subsets were defined as naïve (CD45RO^-^CCR7^+^), CM (CD45RO^+^CCR7^+^), TEMRA (CD45RO^-^ CCR7^-^) and EM (CD45RO^+^CCR7^-^) cells. NK cell subsets include CD56^dim^CD16^+^, CD56^dim^CD16^-^, CD56^bright^CD16^-^ and CD56^bright^CD16^+^ cells. Mean ± SD is shown. **B**. Representative flow cytometry plots and frequencies of CD8^+^ T, CD4^+^ T and NK cells in lung perfusates expressing the makers CD69, CD103, CD49a, PD-1 and CX3CR1 at the indicated combinations (mean ± SD, n=10). Gating strategy is shown in Fig. S3B.

With regards to TRM markers, significant proportions of CD69-expressing T and NK cells were detected in perfusion solutions (**Fig. 4B**). Like donor cells in recipient blood, CD69^+^ T cells did not co-express the TRM markers CD103 and CD49a, indicating a similar TRM-like phenotype as one of the major subsets (**Fig. 4B, Fig. S3B**). Besides CD69, CD103 and CD49a we also analyzed for expression of additional TRM markers by flow cytometry. Of note, the inhibitory receptor PD- 1 showed a different distribution and was not always co-expressed with CD69, implying an independent regulation from the classical TRM markers. The chemokine receptor CX3CR1, another characteristic marker for true-TRM, was poorly expressed on T cells, while in NK cells, it showed an opposite expression pattern compared to CD69. Conclusively, the analysis of lung perfusion solutions allowed us to identify certain T and NK cell subsets with similar phenotypes like donor T and NK cells in recipient blood and, thus, may share their pulmonary origin.

### Lung parenchyma and trachea contain subsets of TRM and TRM-like T and NK cells with different migratory capacities

In order to determine the potential origin of the perfusate, as well as donor T and NK cells found in LTx recipients, we analyzed the lymphocyte composition of human native lung tissue using explanted recipient lung parenchyma and donor trachea tissue as source of pulmonary immune cells (**Table 2**). In lung parenchyma, CD8^+^ and CD4^+^ T cells were present at similar proportions, while NK cells represented 15% of all lymphocytes (**Fig. S4A**). The majority of T cells showed an EM phenotype although TEMRA cells were also highly present in CD8^+^ T cell populations (**Fig. 5A, Fig. S4B**). Only a weak negative correlation between naïve CD4^+^ and CD8^+^ T cell frequencies and recipient age was found in lung parenchyma (**Fig. S4C**). NK cells primarily displayed a CD56^dim^ phenotype, although CD56^bright^ NK cells were present at significant frequencies (**Fig. 5A, Fig. S4B**). As expected, the majority of CD8^+^ T cells in lung parenchyma with ∼60% expressed CD69 and a large proportion co-expressed CD103 and CD49a (**Fig. 5B**). CD69 was present at similar levels in CD4^+^ T cells but with lower co-expression of CD103 and CD49a compared to CD8^+^ T cells. Most CD8^+^ and CD4^+^ T cells expressed PD-1, although a significant proportion expressed this marker independently from CD69. In contrast to T cells, only a minor proportion of lung parenchyma NK cells were CD69^+^ (mean 20%), while CD103 and CD49a were expressed at even lower levels (**Fig. 5B**). Populations of CD69^+^CD103^-^CD49a^-^ TRM-like cells were found among T and NK cells in lung parenchyma and they could, therefore, represent the potential origin of donor as well as perfusate lymphocytes. Based on these subsets, classical CD69^+^CD103^+^CD49a^+^ TRM T cells are supposed to represent truly tissue-resident memory lymphocytes, which seem to be unable to migrate out of the lung into recipient blood or perfusion solution.

**Fig. 5.**
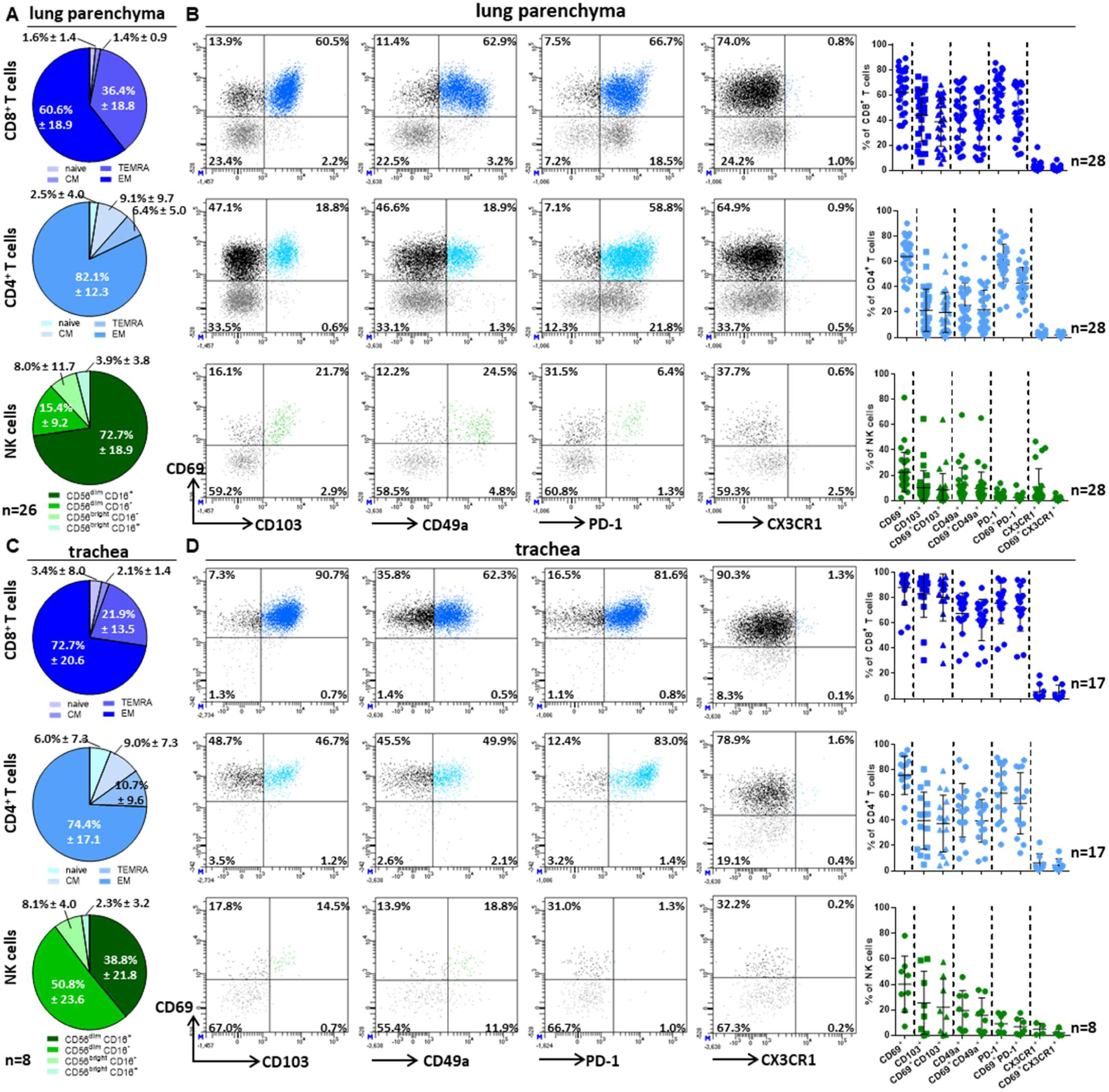
Lung parenchyma and trachea harbor different subsets of TRM cells. **A.** Frequencies of CD8^+^ T cell (n=24), CD4^+^ T cell (n=24) and NK cell (n=26) subsets in lung parenchyma. T cells subsets include naïve (CD45RO^-^ CCR7^+^), CM (CD45RO^+^CCR7^+^), TEMRA (CD45RO^-^CCR7^-^) and EM (CD45RO^+^CCR7^-^) populations. NK cell subsets include CD56^dim^CD16^+^, CD56^dim^CD16^-^, CD56^bright^CD16^-^ and CD56^bright^CD16^+^ populations. Mean ± SD is shown. **B**. Representative flow cytometry plots and frequencies of CD8^+^ T, CD4^+^ T and NK cells (n=23-28 in all groups) in lung parenchyma expressing the makers CD69, CD103, CD49a, PD-1 and CX3CR1 at the indicated combinations. Shown is mean ± SD. **C**. Frequencies of CD8^+^ T cell, CD4^+^ T cell and NK cell subsets in trachea (n=8 in all groups). Mean ± SD is shown. **D**. Representative flow cytometry plots and frequencies of CD8^+^ T cells (n=16- 17), CD4^+^ T cells (n=15-16) and NK cells (n=7-8) in trachea expressing the markers CD69, CD103, CD49a and PD-1 at the indicated combinations. Shown is mean ± SD.

**Table 2.** Demographic characterization of lung tissue (FACS)

| Characteristics | Lung Parenchyma (n=28) | Trachea (n=17) |
| --- | --- | --- |
| **Age at LTx (years)** | 48.6±16.5 |  |
| **Gender** |  |  |
| male | 14 (50.0%) |  |
| female | 14 (50.0%) |  |
| **Transplant indication** |  |  |
| COPD | 14 (50.0%) |  |
| IPF | 6 (21.4%) |  |
| CF | 1 (3.6%) |  |
| PHT | 4 (14.3%) |  |
| ReTx | 2 (7.1%) |  |
| Others | 1 (3.6%) |  |
| **Donor age (years)** |  | 45.9±20.4 |
| **Donor gender** |  |  |
| male |  | 10 (58.8%) |
| female |  | 7 (41.2%) |
| CF: Cystic Fibrosis; COPD: Chronic Obstructive Pulmonary Disease; IPF: Idiopathic Pulmonary Fibrosis; PGD: Primary Graft Dysfunction; PHT: Pulmonary Hypertension; ReTx: Retransplantation<br>Age data are mean values ± standard deviation (SD). |  |  |

In donor trachea tissue, CD8^+^ T cells were overrepresented compared to CD4^+^ T cells, while NK cells were detected at very low frequencies (mean 4.1 %, **Fig. S4D**). In this unique airway tissue, the vast majority of CD8^+^ and CD4^+^ T cells showed an EM phenotype, whereas CD56^dim^CD16^-^ represented the major NK cell population (**Fig. 5C, Fig. S4E**). In the trachea, age did not have any impact on the T cell distribution (**Fig. S4F**). Virtually all CD8^+^ T cells detected in the trachea expressed CD69 and the majority co-expressed the TRM markers CD103, CD49a and PD-1 (**Fig. 5D**). CD69 was detectable at comparable frequencies by CD4^+^ T cells, which preferentially co-expressed CD103 and CD49a albeit at lower levels than CD8^+^ T cells. A proportion of trachea NK cells expressed CD69, CD103 and CD49, but at generally lower frequencies than T cells (**Fig. 5D**). These subset analyses clearly demonstrated that the trachea is highly enriched of classical TRM T and NK cells.

In addition, we directly compared the expression of TRM markers by T and NK cells between the different compartments blood, perfusion solutions, lung parenchyma and trachea and clustered them hierarchically. Interestingly, the composition of T and NK cells in perfusion solutions clustered closer to blood cells than to cells present in lung parenchyma and the trachea (**Fig. 6A- B**). For some markers, such as CD69 and CX3CR1, perfusate cells showed a rather intermediate phenotype between blood and lung tissue (**Fig. 6C, Fig. S5**). Nevertheless, the CD69^+^CD103^-^ CD49a^-^ TRM-like phenotype of perfusate T and NK cells was also found in lung parenchyma (**Fig. 5B**, **Fig. 6C**), indicating that this particular subset may represent the “mobile” tissue-resident memory-like compartment of the lung.

**Fig. 6.**
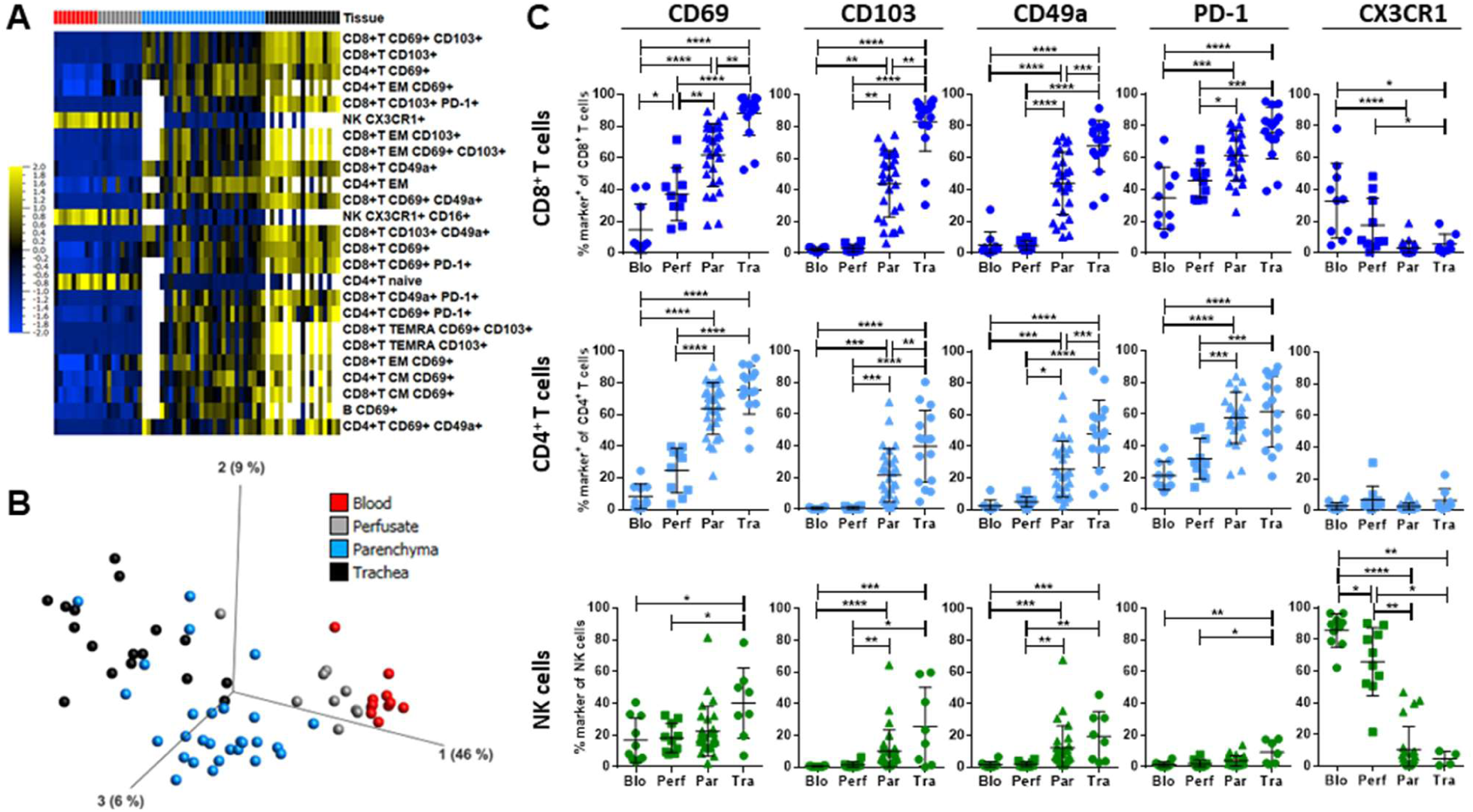
Blood, perfusate, lung parenchyma and trachea lymphocytes express TRM markers at different levels. The lymphocyte composition, with a special focus on markers related to tissue residency, was determined via flow cytometry and compared between blood (Blo, n=10), lung perfusate (Perf, n=10), lung parenchyma (Par, n=43), trachea (Tra, n=17) and lung draining lymph node (n=38) samples. Blood samples correspond to pretransplant blood. **A**. Heatmap generated after performing a multigroup comparison between the different tissues using a panel of 130 immune cell subsets. With a q-value cut-off of 0.05, frequencies of 126 subsets were found significantly different among the groups. Subsets are ordered in the heatmap according to increasing q-values. Shown are only the 30 subsets with the highest q-value. The yellow to blue scale indicates the prevalence of each subset. Missing values are highlighted in green. **B**. Principle component analysis (PCA) performed including the 126 subsets whose frequencies differed significantly between the tissues analyzed. **C**. Proportions of CD8^+^ T cells (upper panel), CD4^+^ T cells (middle panel) and NK cells (lower panel) expressing the markers CD69, CD103, CD49a, PD-1 and CX3CR1 (mean ± SD). Statistical analysis: ANOVA test with Tukey multiple comparison test or Kruskal-Wallis test with Dunn’s multiple comparison test were performed. Asterisks indicate p values (*p<0.05, **p<0.01, ***p<0.001, ****p<0.0001).

### Single cell mRNA transcriptomics identify CD69^+^CD103^+^CD49a^+^ true-TRM and CD69^+^CD103^-^CD49^-^ TRM-like T cells in lung parenchyma

To gain a deeper insight into the T cell compartment of the lung and its molecular characteristics, we performed scRNA sequencing with 60,341 cells isolated from lung parenchyma of 10 patients comprising emphysema (n=5), fibrosis (n=2), pulmonary arterial hypertension (PAH, n=1) and non-malignant (“normal”) tissue of allografts sampled directly before transplantation (n=2; **Table S1**) using the Seq-well technology (20). We clustered 10.799 T cells into 14 biologically relevant sub-clusters (**Fig. 7A, Table S4**) and annotated them as circulating, TRM and TRM-like T cell populations based on the same strategy that we applied for immune phenotyping (**Fig.7B**). We confirmed the definition of T cell subsets in tissue compartments, such as blood with circulatory phenotypes, trachea with enrichment of classical true-TRM and parenchyma with a mixture of TRM and circulatory T cells. Accordingly, we used *CD69* to separate CD69^-^ circulating T cell clusters from CD69^+^ activated or tissue-resident T cell clusters (**Fig. 7B, C**), which resulted in seven circulatory and seven tissue resident/activated T cell clusters (0–13). Apart from cluster 3, all CD69^+^ T cell clusters were grouped close to each other in the upper part of the UMAP, while most of the CD69^-^ clusters were located in the lower part of the UMAP, suggesting that this annotation strategy separated TRM and circulatory T cells. To further separate CD69^+^ activated from CD69^+^ TRM T cells, we incorporated additional T cell activation markers like *HLA-DRA, HLA-DRB, IL2RA* (CD25) (**Fig. 7D**). As had to be expected from TRM T cells, these express *CD69* in conjunction with tissue-residency/retention markers but without activation markers (10). The strongest signal for activation demarcated primarily cluster 3, positioned in the proposed circulatory compartment in the UMAP visualization (**Fig. 7A, D**). Therefore, other clusters in the upper part of the UMAP expressing *CD69^+^ HLA-DRA^-^/DRB^-^ IL2RA^-^* are supposed to represent the TRM compartment. Furthermore, effector and cytotoxic gene signatures showed the strongest signals also in the proposed *CD69^+^HLA-DRA^-^/B^-^ IL2RA^-^* TRM-like clusters (**Fig. 7E**). This is in line with published findings that TRM T cells are able to accumulate mRNA of effector genes (10). Of note, three clusters (clusters 10-12) separated from most other T cell clusters and clusters 10 and 11 displayed higher expression of genes involved in proliferation, i.e. *MKI67* and *TOP2A* (**Fig. 7A, Table S4**), highlighting a discrete profile of proliferating T cells. According to our definition, both proliferating clusters 10 and 11 displayed a circulating phenotype with low *CD69* expression, which again is in line with a lower proliferation capacity of TRM T cells (10). In addition, cluster 12 also showed low *CD69* but higher expression of cell cycle genes like *PTTG1* and *COPS6*. Due to their UMAP proximity to TRM-associated clusters, these cells might to represent a proliferating CD69^lo^ TRM subset.

**Fig. 7.**
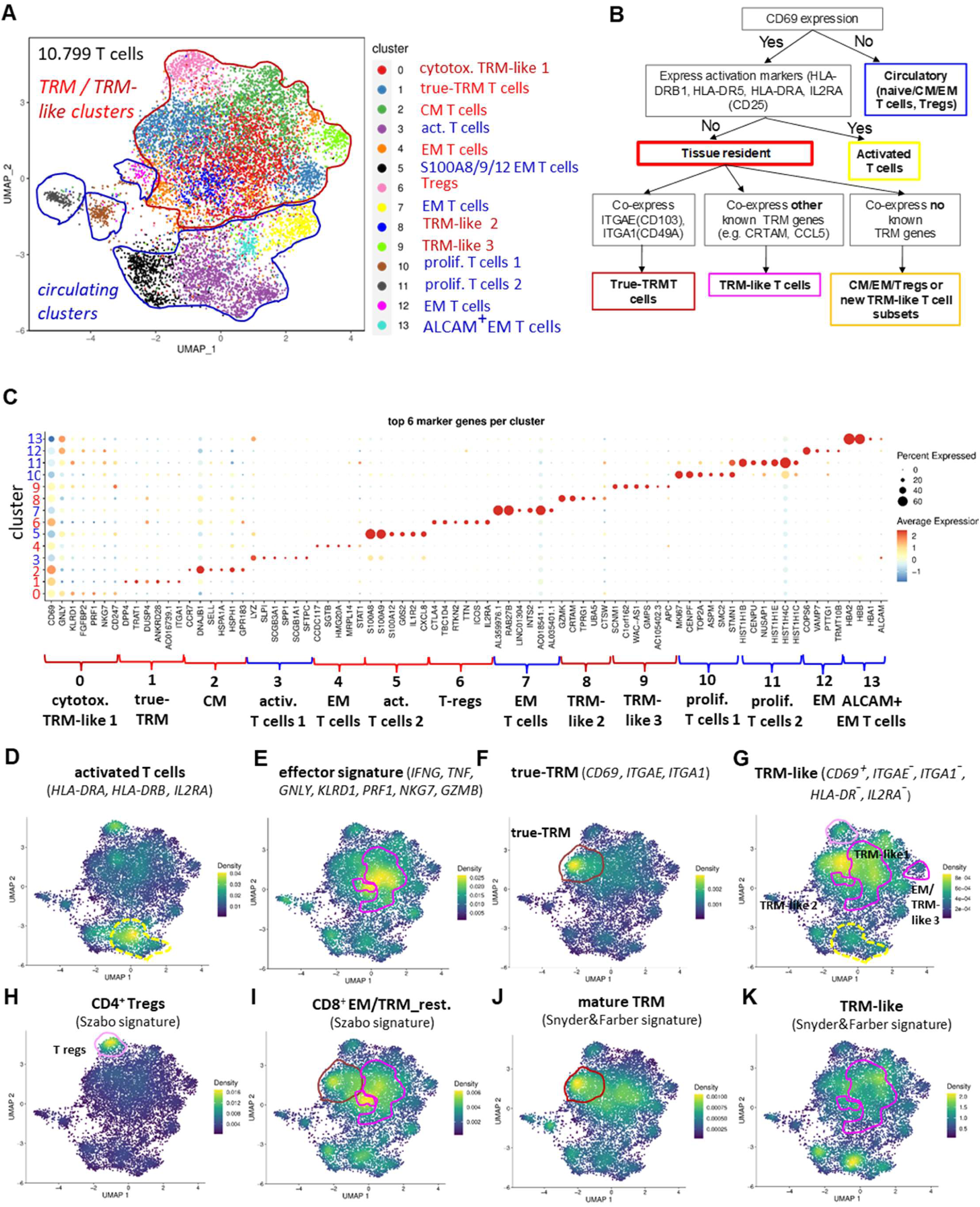
Single cell RNA transcriptomics identifies CD69^+^CD103^-^CD49a^-^ TRM-like and CD69^+^CD103^+^CD49a^+^ true-TRM subsets in lung parenchyma. (A) UMAP dimensionality reduction of lung parenchyma T cells of 10 patients with emphysema (n=5), fibrosis (n=2), PAH (n=1), and Tu. free section of lung tumor patients (n=2). **(B)** Schematic representation of the applied annotation strategy. **(C)** Dot plot showing the top 7 markers per cluster. **(D-K)** UMAP as in (A), clusters expressing a given marker gene signature are indicated. Relative expression is presented as gradient between blue (no expression) and yellow (high expression). Dashed or colored lines indicates clusters as in (A) reflected by color-code. Abbreviations: TRAC: T cell receptor alpha chain constant region; DE = differential expression; UMAP: Uniform Manifold Approximation and Projection for Dimension Reduction.

In addition, a Treg subset, cluster 6, was annotated with enhanced expression of *IL-2RA* (CD25), *CTLA4* and *TIGIT* (**Fig. 7H**) with a highly overlapping signature for CD4^+^ Tregs defined by others (21). Moreover, a naïve/ central memory (CM) T cell cluster 2 was annotated, co-expressing the lymph node homing markers *CCR7* and *SELL* (CD62L) (22). One true-TRM T cell cluster 1 was identified, which co-expressed a signature of CD69 together with the integrins *ITGAE* (CD103) and/or *ITGA1* (CD49a) involved in tissue retention and residency (**Fig. 7F**). This cluster exhibited elevated expression of *RBPJ*, involved in Notch signaling, *CD52*, *IL7R* and several other genes associated with the TRM program (19). In contrast, we found several TRM-like T cell clusters (dark red), which were defined by expression of CD69, but exclusion of activation or true-TRM signatures (**Fig. 7G**). Clusters expressing known TRM-associated genes e.g. *CRTAM, TMSB4X, CCL5, IL32, GZMK, IFNG* were annotated as known TRM-like cells (**Table S4**), while other *CD69^+^* clusters, which did not fit to true-TRM, TRM-like, CM or activated T cells, may represent EM cells or TRM-like subsets with still distinct features. Of note, only a part of cluster 1 showed a true-TRM signature expression, while another part seemed to be more related to TRM-like cells.

This overlap argues for a potential transition from true-TRM to TRM-like subsets and *vice versa* (**Fig. 7A, F, G**). To support our annotation, we also applied published signatures for TRM subsets (19, 21) and could confirm a high overlap between these signatures and our annotations (**Fig. 7I- K**). Taken together, our scRNA sequencing data of human lung parenchyma support our previous findings of two major functionally and phenotypically different tissue-resident memory T cell populations: i) true-TRM T cells co-expressing a set of retention/ residency genes, i.e. *CD69, CD103, CD49a* and others and ii) *CD69^+^CD103^-^CD49a^-^* TRM-like T cells with several sub- clusters defined by distinct signatures. Thus, T cells present in human explanted lung parenchyma represent a “gradient” of true-TRM, TRM-like to circulating T cells, which are likely to be localized in different compartments: true-TRM T cells in close proximity to the airways *vs*. circulating T cells within the interstitial or endothelial compartment. The newly defined TRM-like T cell subsets would represent an intermediate population with migratory capacities and, hence, is likely to resemble donor T cells in recipient blood after LTx.

## Discussion

Allo-recognition by various immune cells represents a clinically relevant obstacle for the success of lung transplantation. Recipient immune cells are able to recognize the transplanted organ as foreign, initiating allo-responses which can ultimately lead to acute or chronic allograft rejection. In parallel, donor lung-resident immune cells transferred into the recipient with the graft may also play a critical role in acceptance or rejection. In this study, we provide new insights on the pulmonary immune system and the interplay between donor and recipient immunity after lung transplantation. In this study, we discovered in a large cohort of 97 lung transplant recipients major dynamics in the lymphocyte composition in recipient blood directly after LTx. These changes have previously been unappreciated, since early time points after transplantation (D0, D1) are not routinely studied, mainly due to logistic requirements of sampling. Despite the heterogeneity of our cohort in terms of genetic background, age, sex and end-stage organ failure, homogeneous changes in lymphocyte composition were detected in all patients, confirming a major and conserved impact of LTx on recipient immunity. Our analysis revealed that T cell subsets, especially CD4^+^ T cells, decreased directly after transplantation accompanied by a relative increase in NK cell frequencies. The presence of passenger donor lymphocytes may contribute to these altered proportions, given that NK cells were the most prominent population among donor cells. However, this explanation is incomplete, since the numbers of all lymphocyte subsets, including NK cells, in blood declined directly after transplantation but to different degrees, with CD4^+^ T cells experiencing the most abrupt decline among all lymphocytes. Extended cell death appears to be an unlikely explanation for this decline, given that live-dead staining did not show increased proportion of dead cells (data nor shown). Therefore, we favor the interpretation that circulating immune cells are relocated to secondary lymphoid organs and / or the transplanted lung, which is, of course, difficult to prove in humans due to the limited access to explanted lung and other tissues. Another explanation could be that the egress of immune cell from peripheral organs or lymphoid tissues is impaired after transplantation, thereby, reducing lymphocyte counts in blood. This phenomenon has been described in human patients after administration of the immunomodulatory drug FTY720, which promotes the down-modulation of S1PR1, thus inhibiting lymphocyte egress from secondary lymphoid organs (23). Whether a similar mechanism may occur after LTx requires further investigation.

In a subcohort (n=44), selected according to the availability of allele-specific mAb, we could detect donor passenger lymphocytes in all patients, confirming previous findings (17). Our subset analyses identified NK and T cells as major donor subsets, whereas B cells and myeloid cells were almost absent (data not shown). Donor T cells disappeared in recipient blood almost completely three weeks after LTx, which may explain why other publications starting their analyses at later time points failed to detect them (19). In contrast, donor NK cells were still detectable in recipient blood at discharge, i.e. three weeks after transplantation, which argues for either a prolonged release of NK cells or a higher survival and/or proliferation rate in recipient periphery. In terms of their clinical relevance, the frequencies of donor T and NK cells did not differ between lung recipients with or without PGD, indicating that other mechanisms like ischemia/reperfusion injury are primarily responsible for early graft function. However, significantly higher frequencies of donor T cells were characteristic for CLAD-free recipients two years after LTx compared to patients suffering from CLAD, indicating that this transient chimerism especially of donor T cells may be protective for CLAD development. The potential protective mechanisms need to be further investigated but the observation of an higher IFN-γ expression in donor T cells points towards a pre-activation during the *ex vivo* phase, which may enable these cells to partially control the development of allo-reactive T cells.

With regards to naïve or memory subsets, donor passenger T cells were enriched for CM and EM T cells, whereas naïve T cell frequencies were reduced compared to recipient T cells. In addition, donor T and NK cells displayed higher levels of CD69 than recipient cells but lacked expression of TRM markers CD103 and CD49a as well as activation markers CD25 and HLA-DR. Since this peculiar phenotype was also found in perfusate T and NK cells, we argue that these cells do not represent circulating but rather CD69^+^CD103^-^CD49a^-^ TRM-like subsets, as defined in earlier publications in lung tissue (19). Indeed, our scRNA sequencing analyses confirmed the detection of *CD69^+^CD103^-^CD49a^-^* TRM-like subsets in lung parenchyma, with strong accumulation of transcripts associated with cytotoxicity, e.g. *IFNG, GZMB, GNLY*. This argues for an intermediate subset of *CD69^+^CD103^-^CD49a^-^* TRM-like T and NK cells in the lung, which seems to be able to leave the organ after transplantation or into perfusates as ‘mobile TRM-like cells’. ScRNA sequencing was used to confirm and extend our observations on distinct TRM- and TRM-like subsets at the transcriptional level, which were initially defined by the *CD69, CD103, CD49a* and *PD-1* marker set. Of note, our unsupervised clusters of circulating, TRM and TRM-like T cell subsets in lung parenchyma were consistent with transcriptional signatures of these T cell subsets in recent publications (19, 21), which further supports the concept of different lung-resident memory T cell subsets, characterized by activation and/ or effector functions like cytotoxicity. Recently, recipient CD8^+^ T cells with TRM-features and clonal expansion were shown to be associated with acute cellular rejection, suggesting an infiltration and differentiation of selected recipient T cell clones with allo-reactive potential (24).

Of note, TRM and TRM-like T cells were also detected in lung perfusates, which represent a valuable non-invasive source of pulmonary immune cells with potential migratory capacity. These cells are unlikely to represent residual circulating cells, since before explantation, lungs are flushed twice with 4 L perfusion solution, thus, eliminating contaminating donor blood. The finding of significantly enriched proportions of CD69^+^CD103^-^CD49a^-^ CD8^+^ and CD4^+^ T cells, therefore, further supports our interpretation of a common origin of TRM-like T cells in the mobile compartment of the lung, which could be proven by T cell receptor (TCR) sequencing. Further indirect proof is provided by the trachea as pure airway compartment without endothelial or interstitial cells. Since almost all T cells detected in trachea expressed CD69 in combination with the TRM markers CD103, CD49a and PD-1, a localization of TRM T cells in this airway compartment, presumably via CD103/E-Cadherin interactions, can be assumed.

For the skin, it has been demonstrated that cutaneous CD4^+^CD103^+^ TRM T cells that downregulate CD69 have the capacity to leave the dermis and reenter the circulation, also opening up the concept of strict residency of TRM cells towards inflammation-driven mobilization (25). Based on the distinct composition of T cells in trachea, lung parenchyma, perfusates and blood, we propose at least three major T cell subpopulations in human lung parenchyma with transcriptionally distinct subsets: CD69^-^ circulating, CD69^+^CD103^-^CD49a^-^ TRM-like “mobile” and CD69^+^CD103^+^ CD49a^+^ TRM “sessile” T cells.

The distribution of the major NK cell subsets is also remarkable since CD56^dim^ NK cells represented the major subset in donor passenger as well as perfusate NK cells with only a minor fraction of CD56^bright^ NK cells. Depending on the source of lung tissue, tissue-resident NK cells were described as CD56^bright^CD16^-^ NK cells expressing CD69, CD103 and CD49a (14), but also CD56^dim^CD69^+^ NK cells were detected in lung parenchyma (16), which may be related to the variability on tertiary lymphoid structures as part of the lung tissue with its known enrichment of CD56^bright^ NK cells. In our study, primarily CD56^dim^ NK cells were identified in recipient blood, perfusates and lung parenchyma arguing for their substantial contribution to the human lung compartment with both mobile and resident features.

The clinical relevance of donor T cells has been recently shown in lung recipients more than four weeks after LTx: patients with persistent donor TRM T cells in BAL experienced reduced PGD and acute cellular rejection, arguing for a protective function in the airway compartment (19). One limitation of our study is therefore, that complementary BAL samples were not available for identification of donor cells. However, our finding that the frequencies of donor T and NK cells did not differ between PDG and PDG-free recipients, even at the exact time point of PDG assessment during the first 72 h after LTx, underlines the relevance of the different compartments, especially blood and BAL. Regarding chronic allograft dysfunction, we provide evidence of a potential protective effect of donor passenger T cells during first three weeks following LTx since CLAD-free patients two years after LTx seems to have higher donor T cell frequencies compared to CLAD patients. Unfortunately, the rather limited patient number in our CLAD group (n=7) did not allow to draw definite conclusions, which represents another limitation of our study. Further limitations are the single-center design and incomplete availability of HLA-allele specific monoclonal antibodies to analyze all donor HLA combinations.

Taken together and to the best of our knowledge, our study is the first to identify a transient chimerism of donor T and NK cells circulating in recipient blood, starting immediately after LTx and persisting for at least the first month. These donor T and NK cells are characterized by a TRM-like phenotype, which could be found in lung parenchyma, trachea as well as lung perfusates both via flow cytometry and scRNA sequencing. These TRM-like T cells may contribute to a better long-term outcome after lung transplantation although the underlying mechanisms need to be elucidated further.

## Supporting information

Supplementary Material Bellmas-Sanz et al BioRxiv 2023

## Author contributions

RBS performed experiments, analyzed data and wrote major parts of the manuscript. AMH, KAB, JK, KB and LMH supported experiments. BW coordinated sample collection. EC performed experiments, analyzed data and wrote the manuscript. JFK performed experiments and wrote the manuscript. FI, WS, AH and GW performed LTx and assisted with sample collection. KH, MB, TSK and KB performed scRNA sequencing analyses. DJ provided tissue samples and helped writing the manuscript. MG assisted with sample collection. JLS designed and supervised scRNA sequencing experiments. CSF supervised the work, designed experiments, analyzed data and wrote the manuscript.

## Acknowledgements

This project was supported by the German Research Foundation DFG, SFB 738 project B3, FA-483/1-1, JO 743/3-1, SCHU 950/9-1, WA 1700/5-1, WI 4088/3-1and FOR2830 project P07 and, the German Center for Infection Research DZIF TTU-IICH 07.822, and the MHH Transplant Center CORE100 projects 19_09 and 21_06.

---

**Table S1.** Demographic characterization of lung tissue (scRNA sequencing)

| Demographic characterization of lung tissue (scRNA seq.) |  |
| --- | --- |
| Lung Parenchyma (n=16) |  |
| **Characteristics** |  |
| **Age (years)** | 55.4±10 |
| **Gender** |  |
| male | 14 (62.5%) |
| female | 14 (37.5%) |
| **End stage disease/ tu.free tissue** |  |
| Emphysema/ COPD | 6 (38.0%) |
| IPF | 6 (38.0%) |
| PAH/ PHT | 1 (6.0%) |
| Others (tu.free) | 3 (19%) |
| COPD: Chronic Obstructive Pulmonary Disease; IPF: Idiopathic Pulmonary Fibrosis; PGD: Primary Graft Dysfunction; PHT: Pulmonary Hypertension;<br>Age data are mean values ± standard deviation (SD). |  |

**Table S2.**
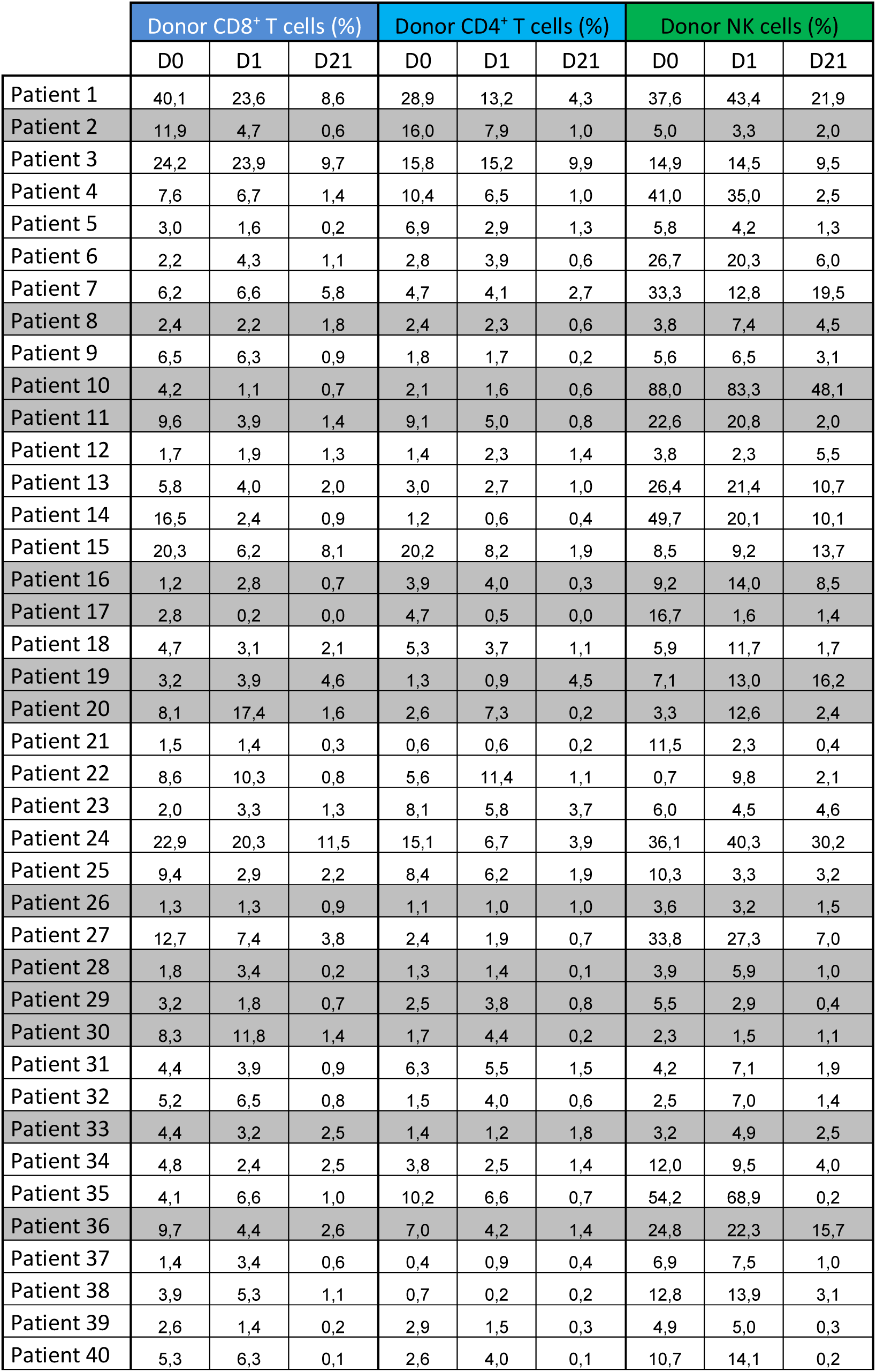
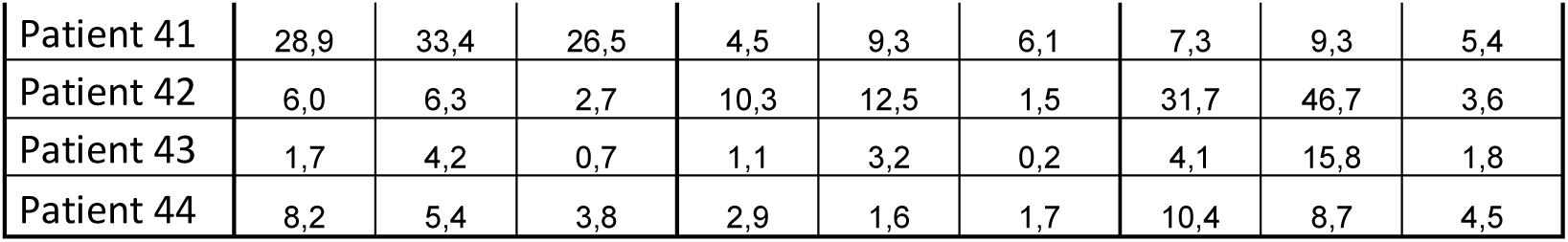
Frequencies of donor T and NK cells.

|  | Donor CD8<sup>+</sup> T cells (%) |  |  | Donor CD4<sup>+</sup> T cells (%) |  |  | Donor NK cells (%) |  |  |
| --- | --- | --- | --- | --- | --- | --- | --- | --- | --- |
|  | D0 | D1 | D21 | D0 | D1 | D21 | D0 | D1 | D21 |
| Patient 1 | 40,1 | 23,6 | 8,6 | 28,9 | 13,2 | 4,3 | 37,6 | 43,4 | 21,9 |
| Patient 2 | 11,9 | 4,7 | 0,6 | 16,0 | 7,9 | 1,0 | 5,0 | 3,3 | 2,0 |
| Patient 3 | 24,2 | 23,9 | 9,7 | 15,8 | 15,2 | 9,9 | 14,9 | 14,5 | 9,5 |
| Patient 4 | 7,6 | 6,7 | 1,4 | 10,4 | 6,5 | 1,0 | 41,0 | 35,0 | 2,5 |
| Patient 5 | 3,0 | 1,6 | 0,2 | 6,9 | 2,9 | 1,3 | 5,8 | 4,2 | 1,3 |
| Patient 6 | 2,2 | 4,3 | 1,1 | 2,8 | 3,9 | 0,6 | 26,7 | 20,3 | 6,0 |
| Patient 7 | 6,2 | 6,6 | 5,8 | 4,7 | 4,1 | 2,7 | 33,3 | 12,8 | 19,5 |
| Patient 8 | 2,4 | 2,2 | 1,8 | 2,4 | 2,3 | 0,6 | 3,8 | 7,4 | 4,5 |
| Patient 9 | 6,5 | 6,3 | 0,9 | 1,8 | 1,7 | 0,2 | 5,6 | 6,5 | 3,1 |
| Patient 10 | 4,2 | 1,1 | 0,7 | 2,1 | 1,6 | 0,6 | 88,0 | 83,3 | 48,1 |
| Patient 11 | 9,6 | 3,9 | 1,4 | 9,1 | 5,0 | 0,8 | 22,6 | 20,8 | 2,0 |
| Patient 12 | 1,7 | 1,9 | 1,3 | 1,4 | 2,3 | 1,4 | 3,8 | 2,3 | 5,5 |
| Patient 13 | 5,8 | 4,0 | 2,0 | 3,0 | 2,7 | 1,0 | 26,4 | 21,4 | 10,7 |
| Patient 14 | 16,5 | 2,4 | 0,9 | 1,2 | 0,6 | 0,4 | 49,7 | 20,1 | 10,1 |
| Patient 15 | 20,3 | 6,2 | 8,1 | 20,2 | 8,2 | 1,9 | 8,5 | 9,2 | 13,7 |
| Patient 16 | 1,2 | 2,8 | 0,7 | 3,9 | 4,0 | 0,3 | 9,2 | 14,0 | 8,5 |
| Patient 17 | 2,8 | 0,2 | 0,0 | 4,7 | 0,5 | 0,0 | 16,7 | 1,6 | 1,4 |
| Patient 18 | 4,7 | 3,1 | 2,1 | 5,3 | 3,7 | 1,1 | 5,9 | 11,7 | 1,7 |
| Patient 19 | 3,2 | 3,9 | 4,6 | 1,3 | 0,9 | 4,5 | 7,1 | 13,0 | 16,2 |
| Patient 20 | 8,1 | 17,4 | 1,6 | 2,6 | 7,3 | 0,2 | 3,3 | 12,6 | 2,4 |
| Patient 21 | 1,5 | 1,4 | 0,3 | 0,6 | 0,6 | 0,2 | 11,5 | 2,3 | 0,4 |
| Patient 22 | 8,6 | 10,3 | 0,8 | 5,6 | 11,4 | 1,1 | 0,7 | 9,8 | 2,1 |
| Patient 23 | 2,0 | 3,3 | 1,3 | 8,1 | 5,8 | 3,7 | 6,0 | 4,5 | 4,6 |
| Patient 24 | 22,9 | 20,3 | 11,5 | 15,1 | 6,7 | 3,9 | 36,1 | 40,3 | 30,2 |
| Patient 25 | 9,4 | 2,9 | 2,2 | 8,4 | 6,2 | 1,9 | 10,3 | 3,3 | 3,2 |
| Patient 26 | 1,3 | 1,3 | 0,9 | 1,1 | 1,0 | 1,0 | 3,6 | 3,2 | 1,5 |
| Patient 27 | 12,7 | 7,4 | 3,8 | 2,4 | 1,9 | 0,7 | 33,8 | 27,3 | 7,0 |
| Patient 28 | 1,8 | 3,4 | 0,2 | 1,3 | 1,4 | 0,1 | 3,9 | 5,9 | 1,0 |
| Patient 29 | 3,2 | 1,8 | 0,7 | 2,5 | 3,8 | 0,8 | 5,5 | 2,9 | 0,4 |
| Patient 30 | 8,3 | 11,8 | 1,4 | 1,7 | 4,4 | 0,2 | 2,3 | 1,5 | 1,1 |
| Patient 31 | 4,4 | 3,9 | 0,9 | 6,3 | 5,5 | 1,5 | 4,2 | 7,1 | 1,9 |
| Patient 32 | 5,2 | 6,5 | 0,8 | 1,5 | 4,0 | 0,6 | 2,5 | 7,0 | 1,4 |
| Patient 33 | 4,4 | 3,2 | 2,5 | 1,4 | 1,2 | 1,8 | 3,2 | 4,9 | 2,5 |
| Patient 34 | 4,8 | 2,4 | 2,5 | 3,8 | 2,5 | 1,4 | 12,0 | 9,5 | 4,0 |
| Patient 35 | 4,1 | 6,6 | 1,0 | 10,2 | 6,6 | 0,7 | 54,2 | 68,9 | 0,2 |
| Patient 36 | 9,7 | 4,4 | 2,6 | 7,0 | 4,2 | 1,4 | 24,8 | 22,3 | 15,7 |
| Patient 37 | 1,4 | 3,4 | 0,6 | 0,4 | 0,9 | 0,4 | 6,9 | 7,5 | 1,0 |
| Patient 38 | 3,9 | 5,3 | 1,1 | 0,7 | 0,2 | 0,2 | 12,8 | 13,9 | 3,1 |
| Patient 39 | 2,6 | 1,4 | 0,2 | 2,9 | 1,5 | 0,3 | 4,9 | 5,0 | 0,3 |
| Patient 40 | 5,3 | 6,3 | 0,1 | 2,6 | 4,0 | 0,1 | 10,7 | 14,1 | 0,2 |
| Patient 41 | 28,9 | 33,4 | 26,5 | 4,5 | 9,3 | 6,1 | 7,3 | 9,3 | 5,4 |
| Patient 42 | 6,0 | 6,3 | 2,7 | 10,3 | 12,5 | 1,5 | 31,7 | 46,7 | 3,6 |
| Patient 43 | 1,7 | 4,2 | 0,7 | 1,1 | 3,2 | 0,2 | 4,1 | 15,8 | 1,8 |
| Patient 44 | 8,2 | 5,4 | 3,8 | 2,9 | 1,6 | 1,7 | 10,4 | 8,7 | 4,5 |

**Tab. S3.**
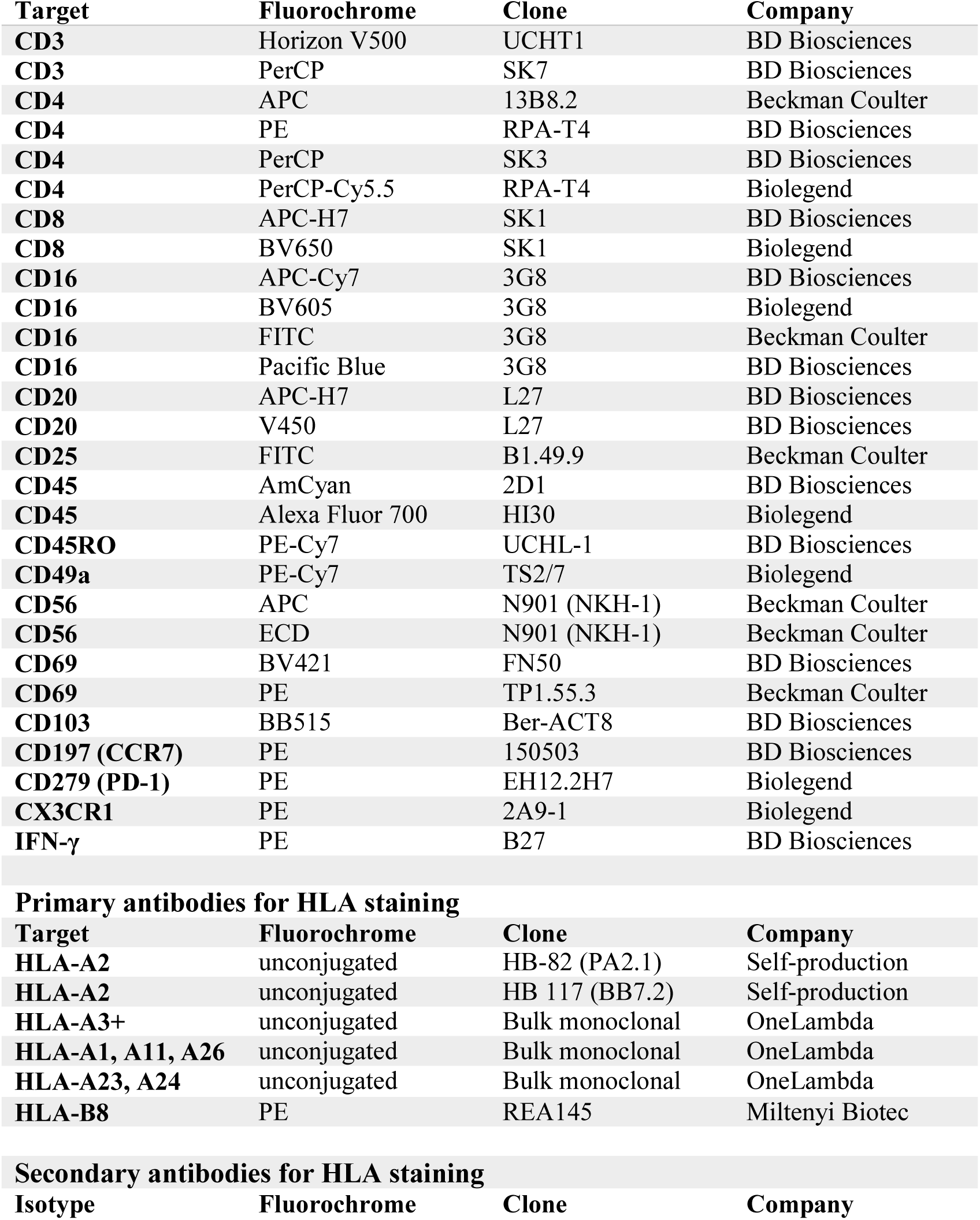
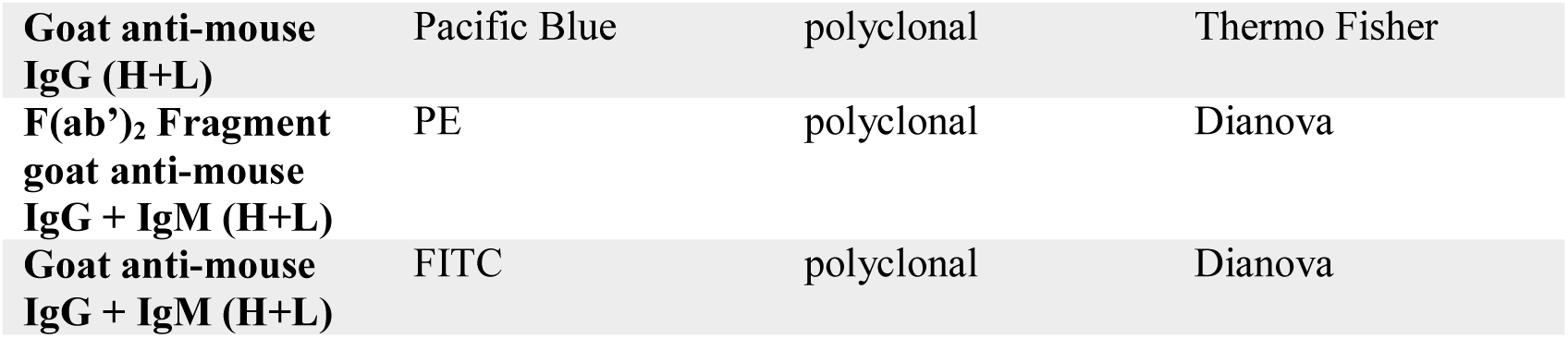
List of antibodies used for flow cytometric analyses.

**Tab. S4.**
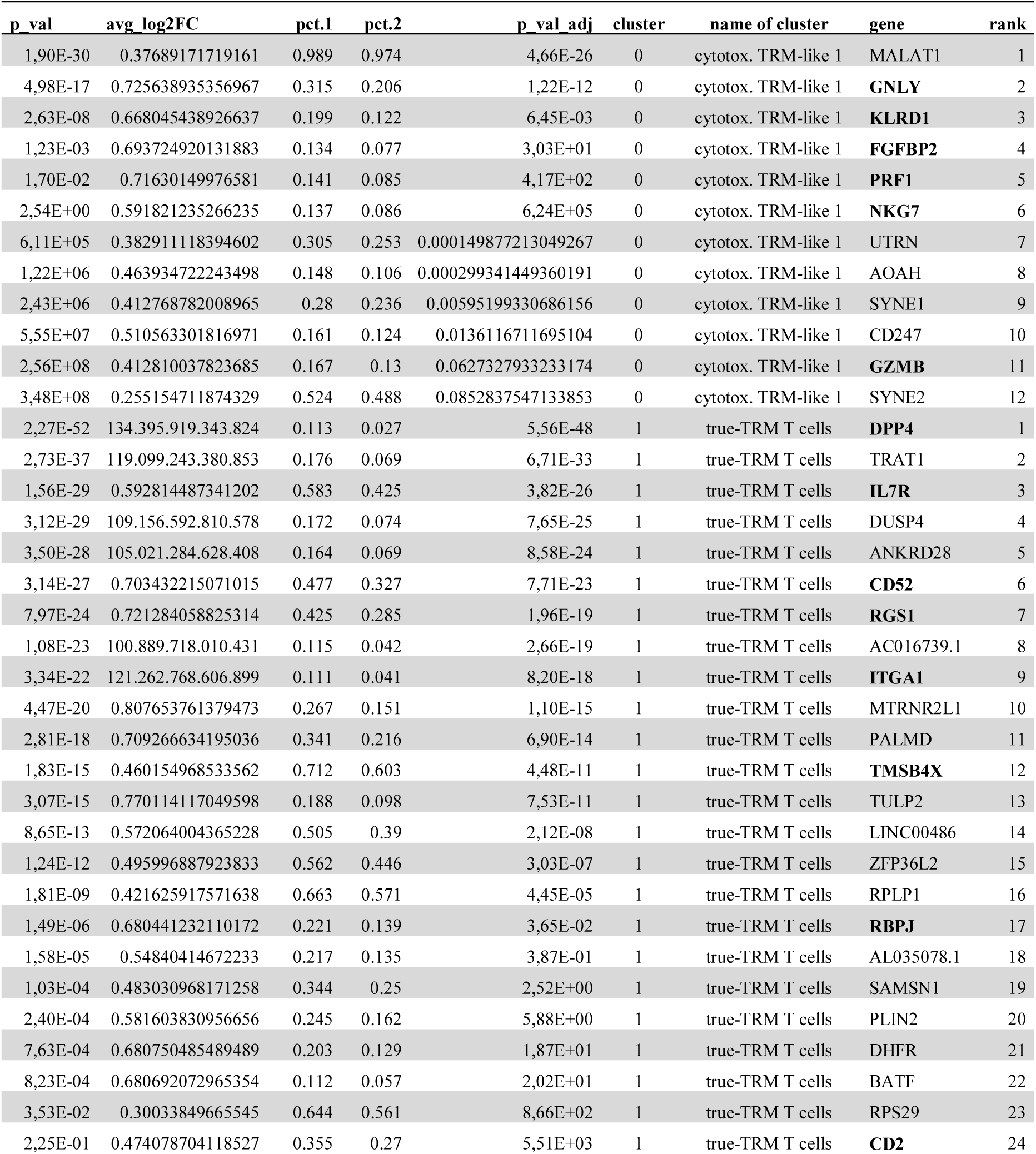
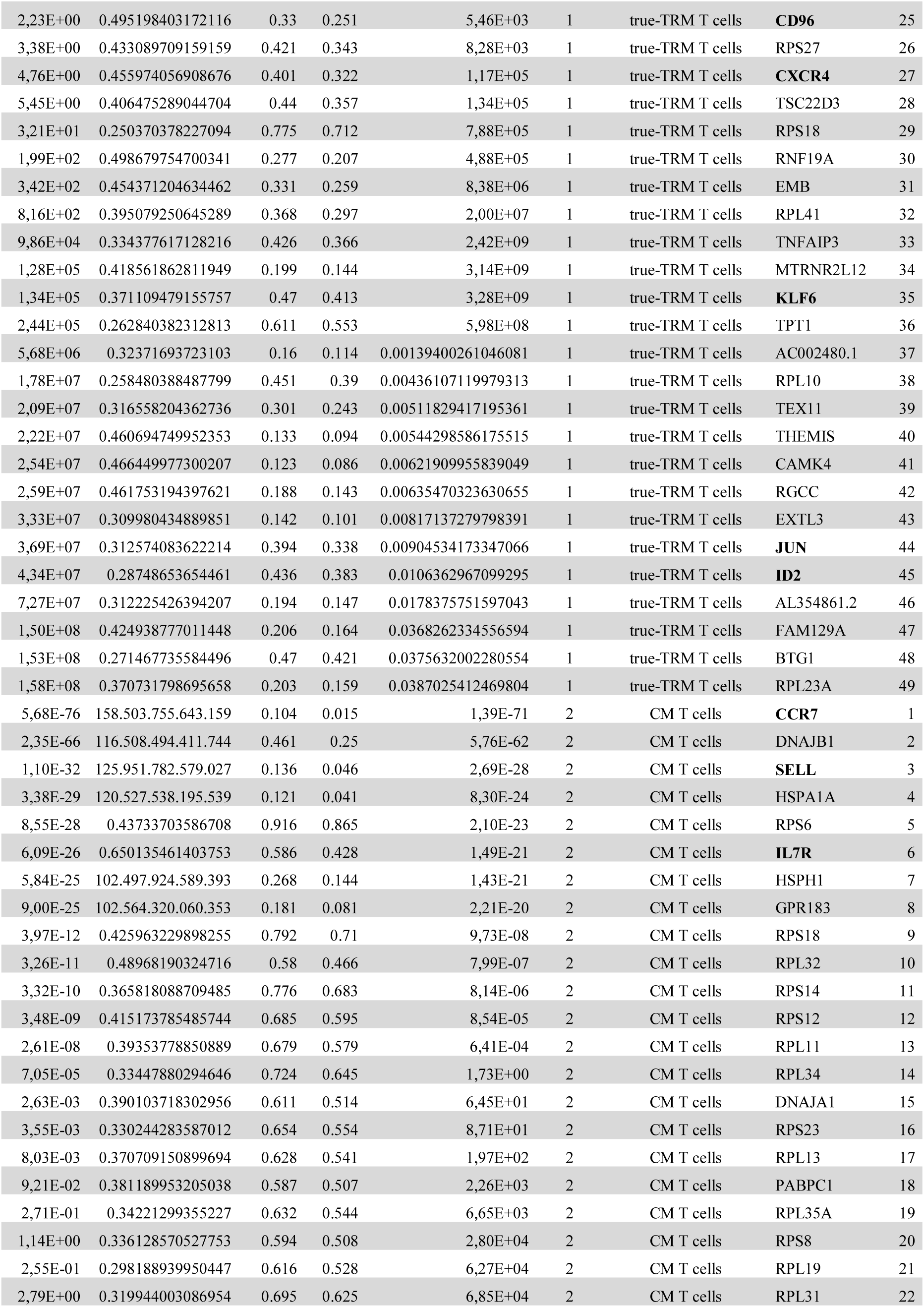
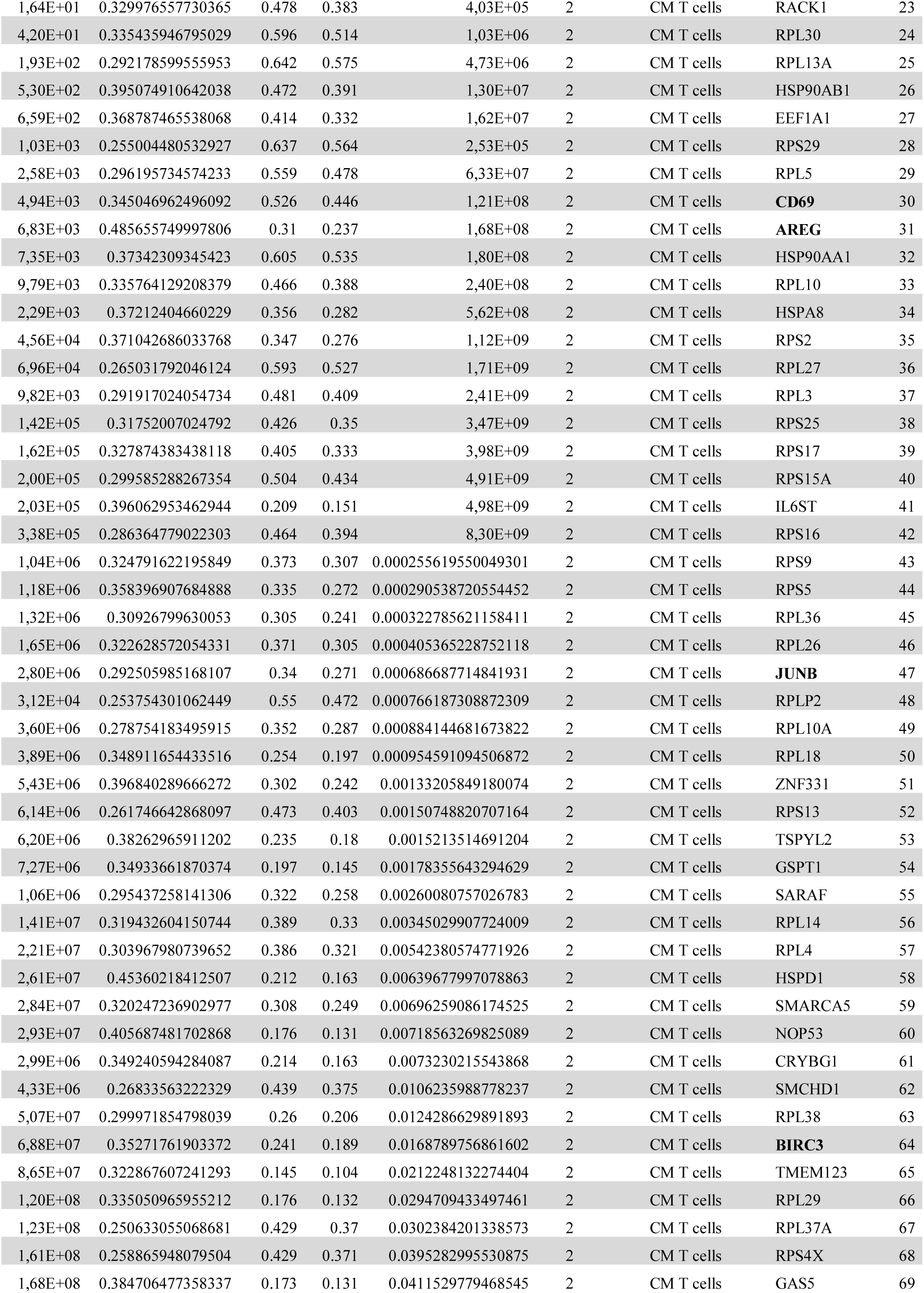
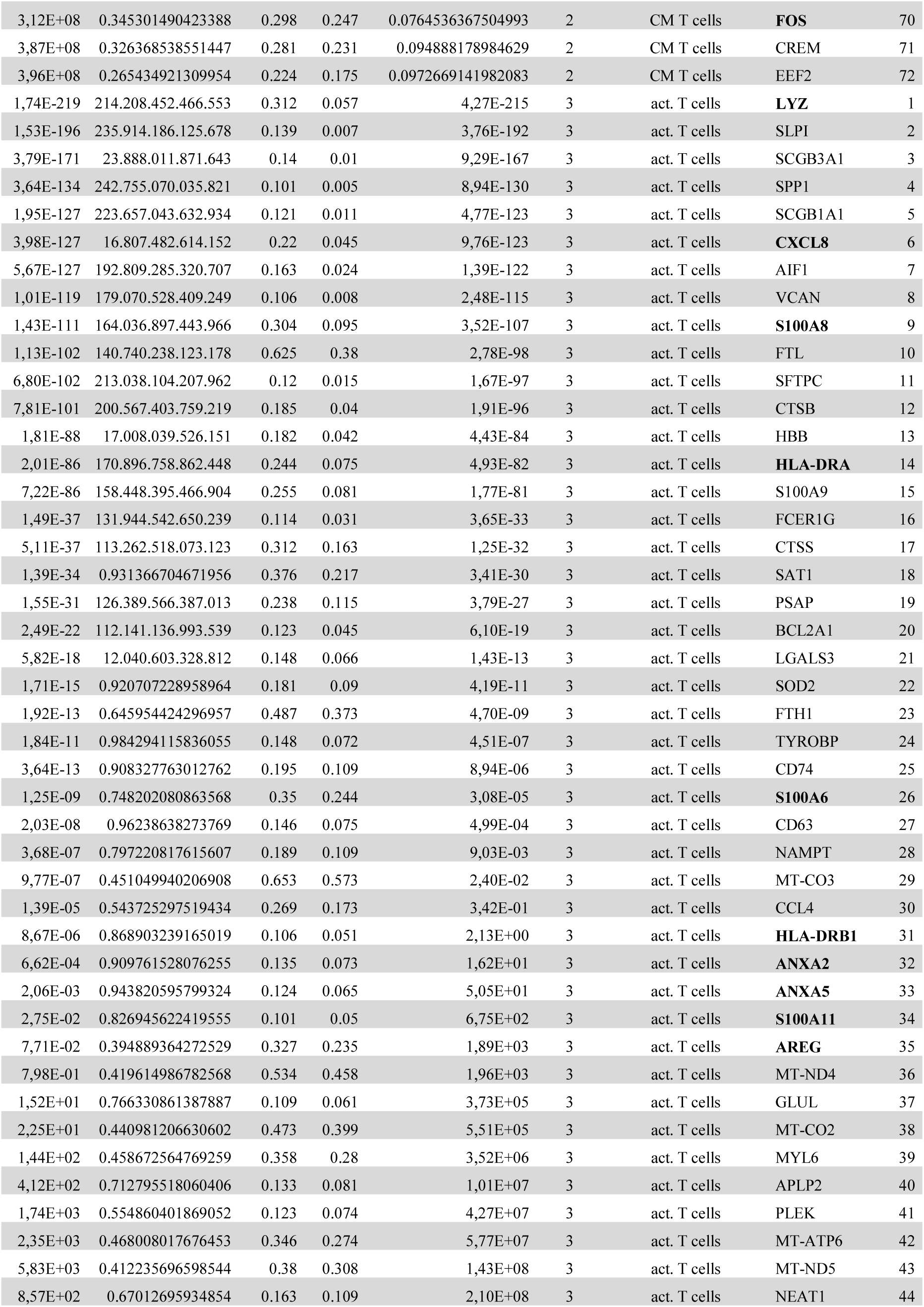
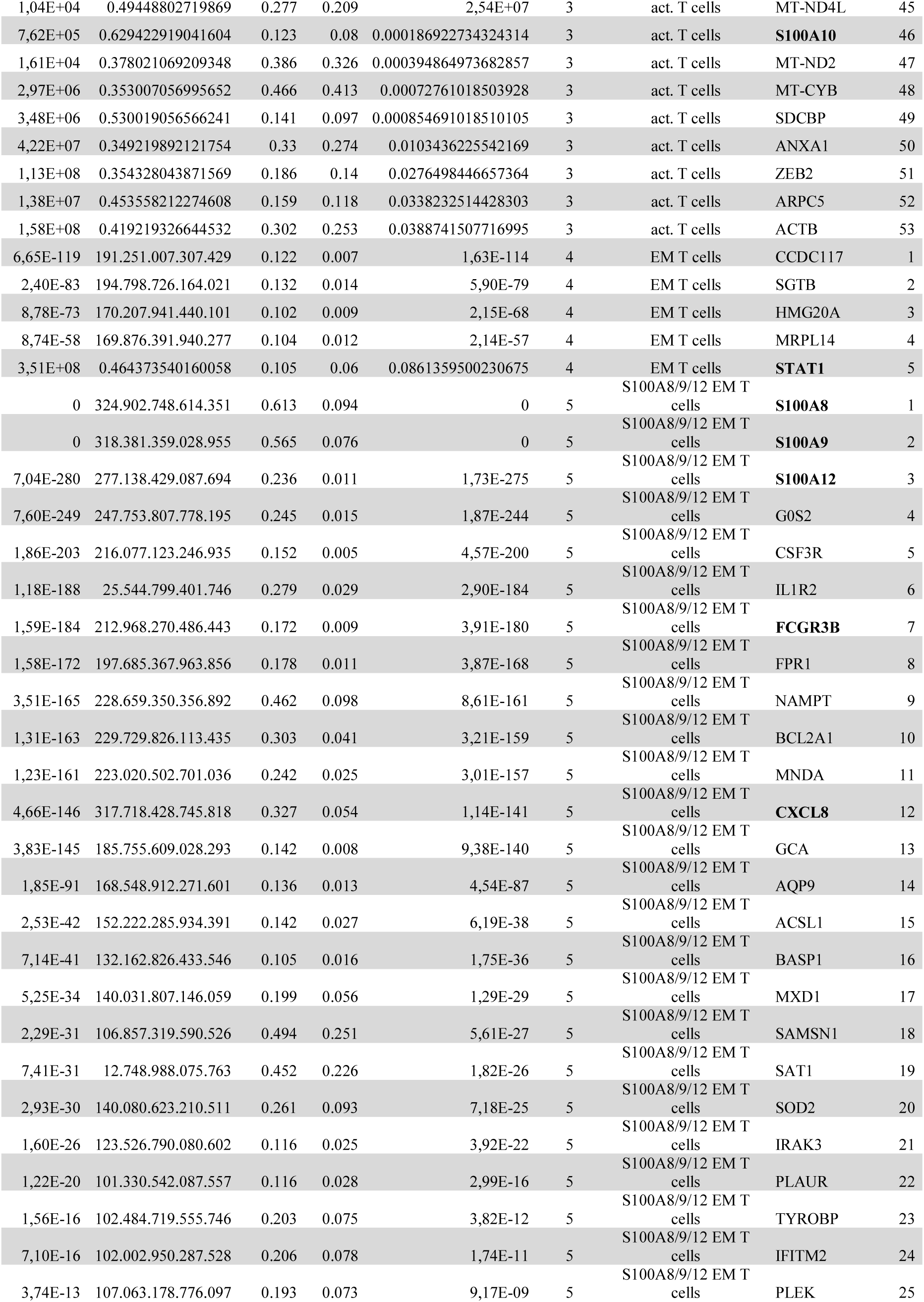
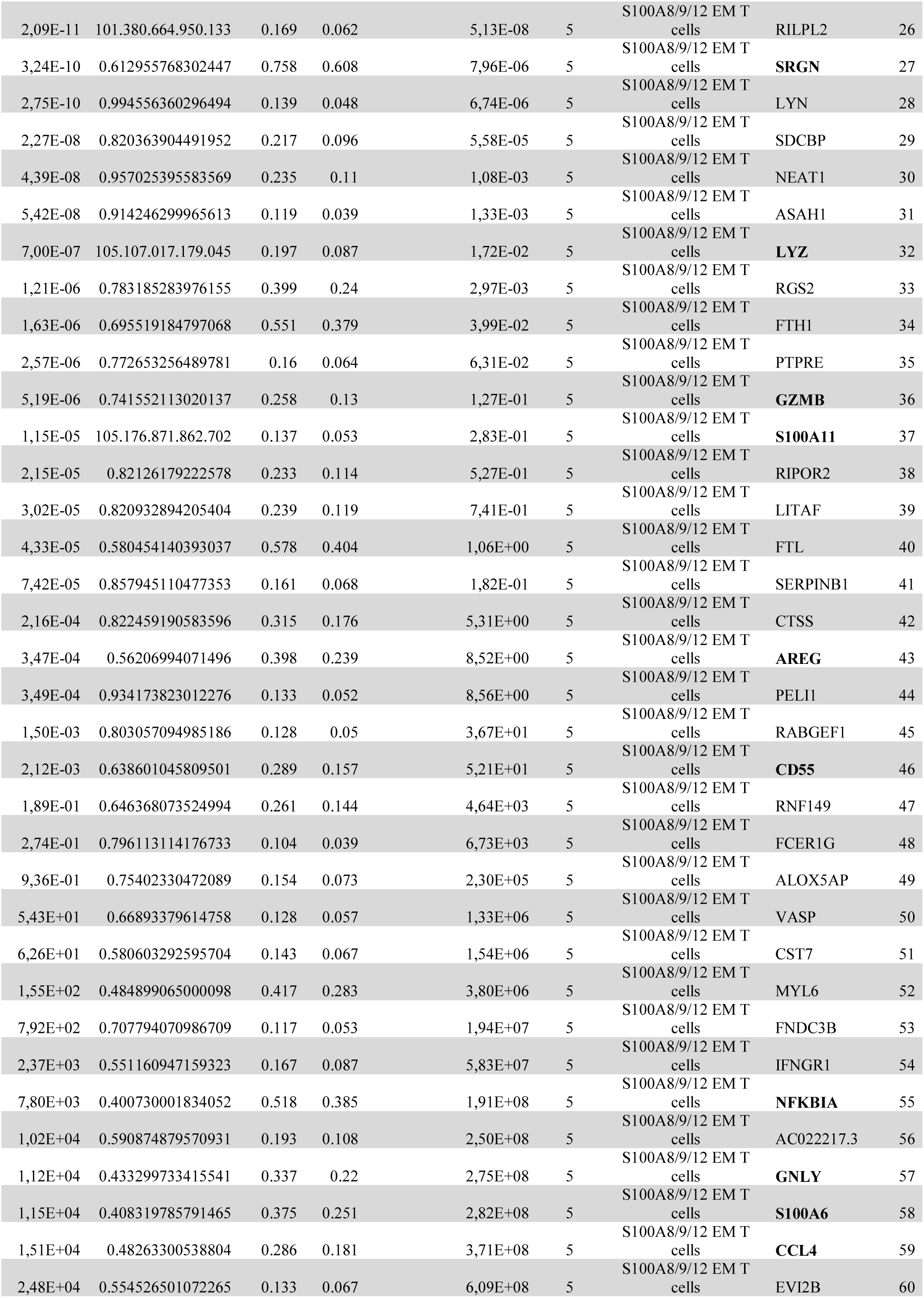
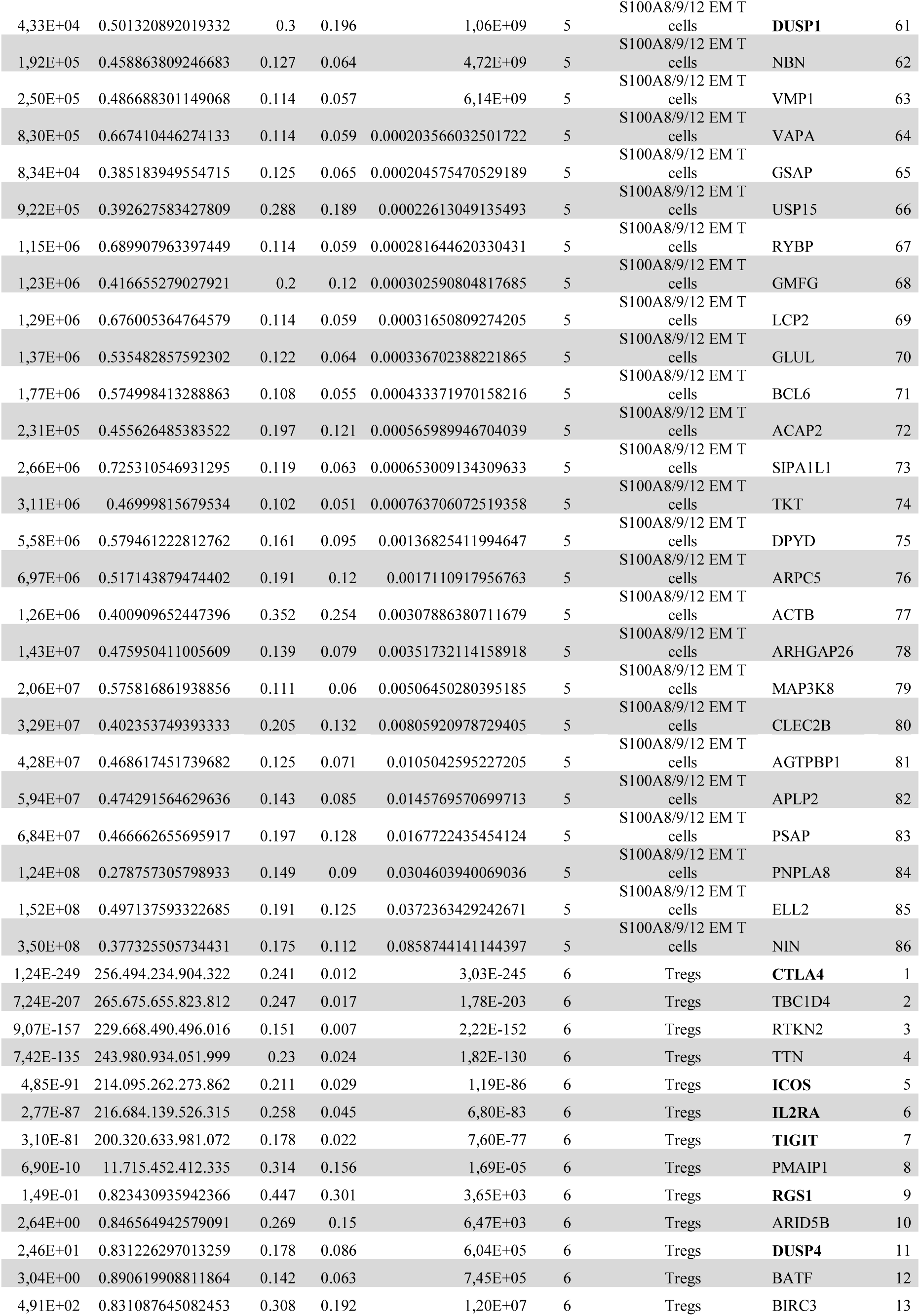
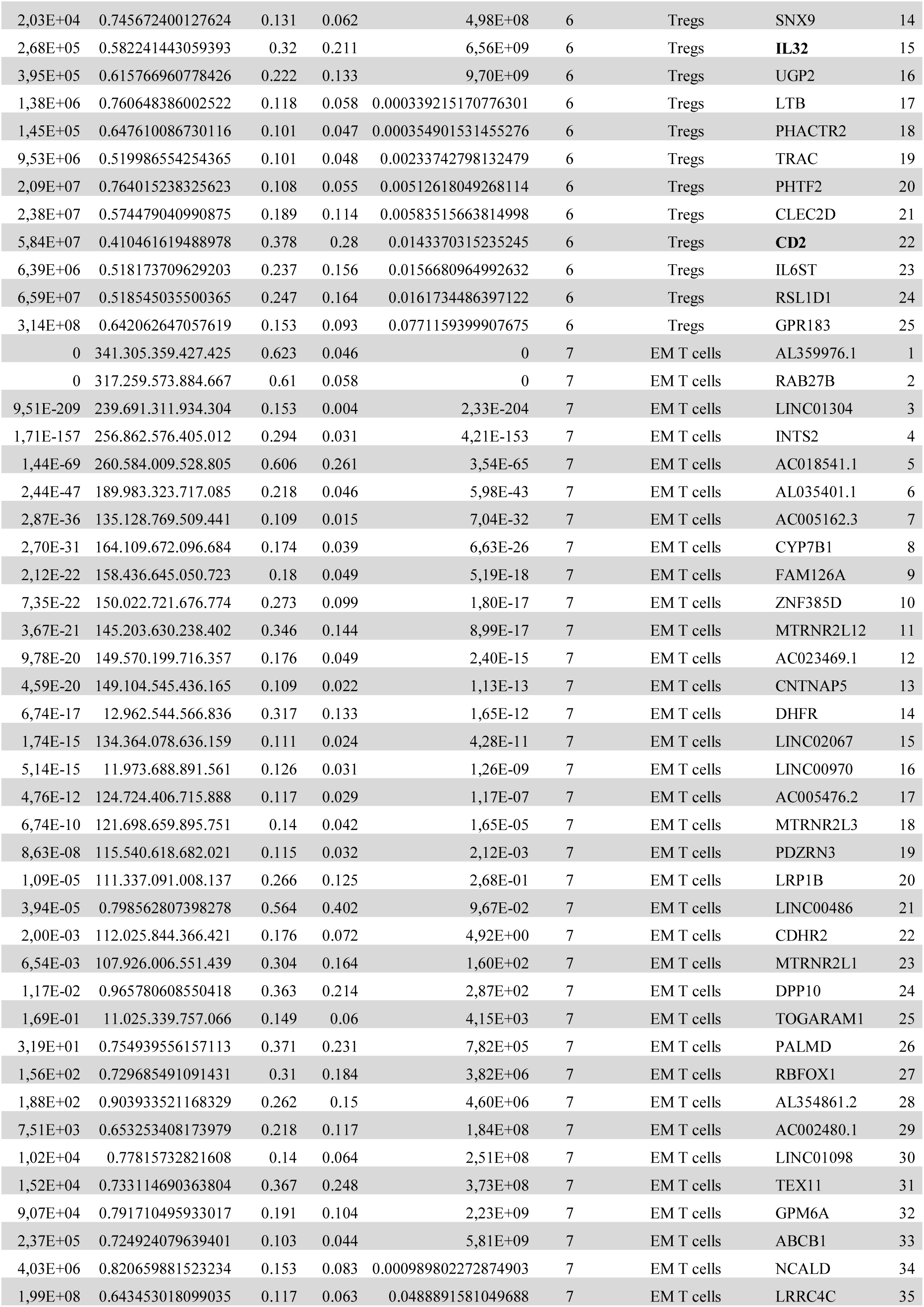
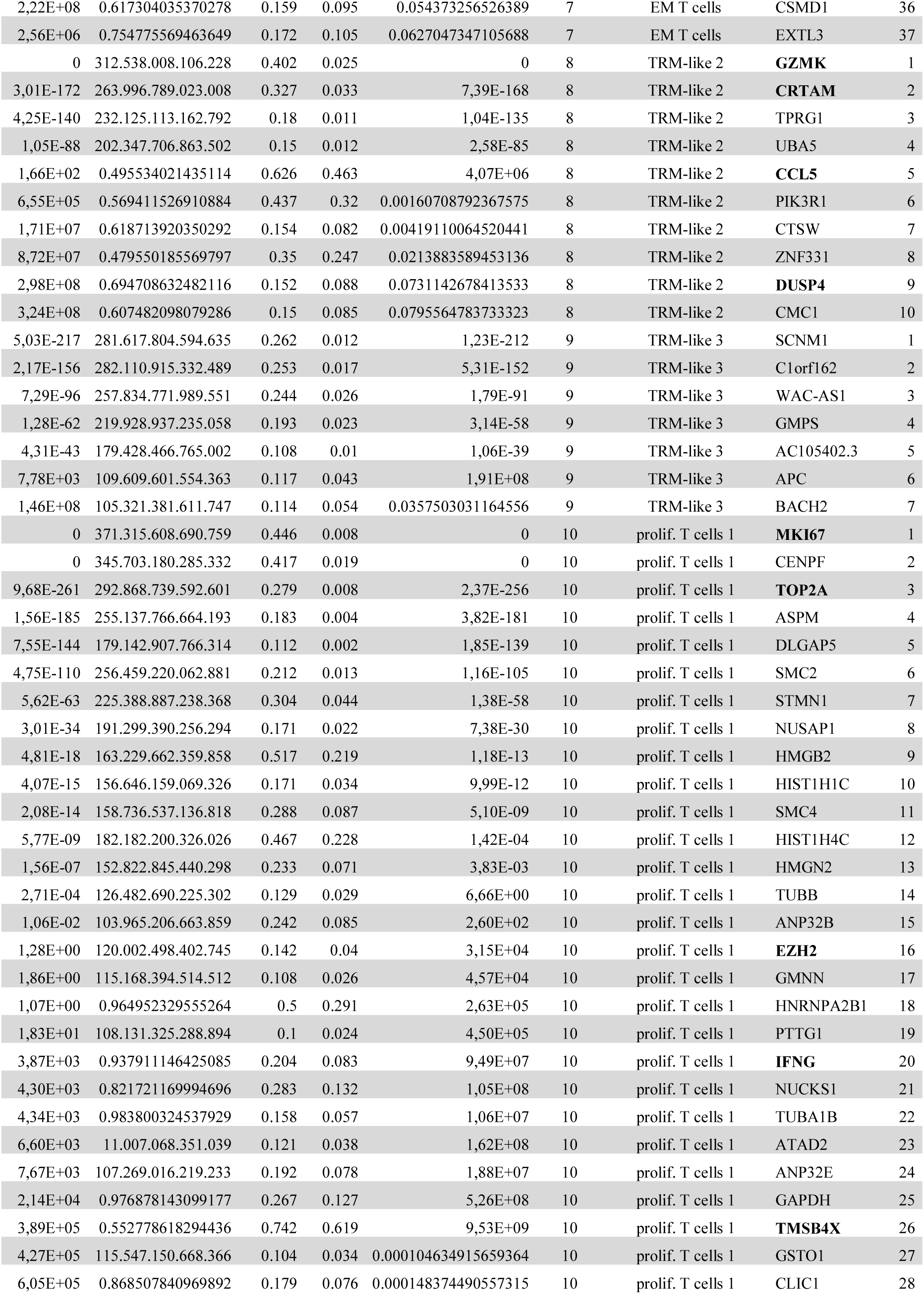
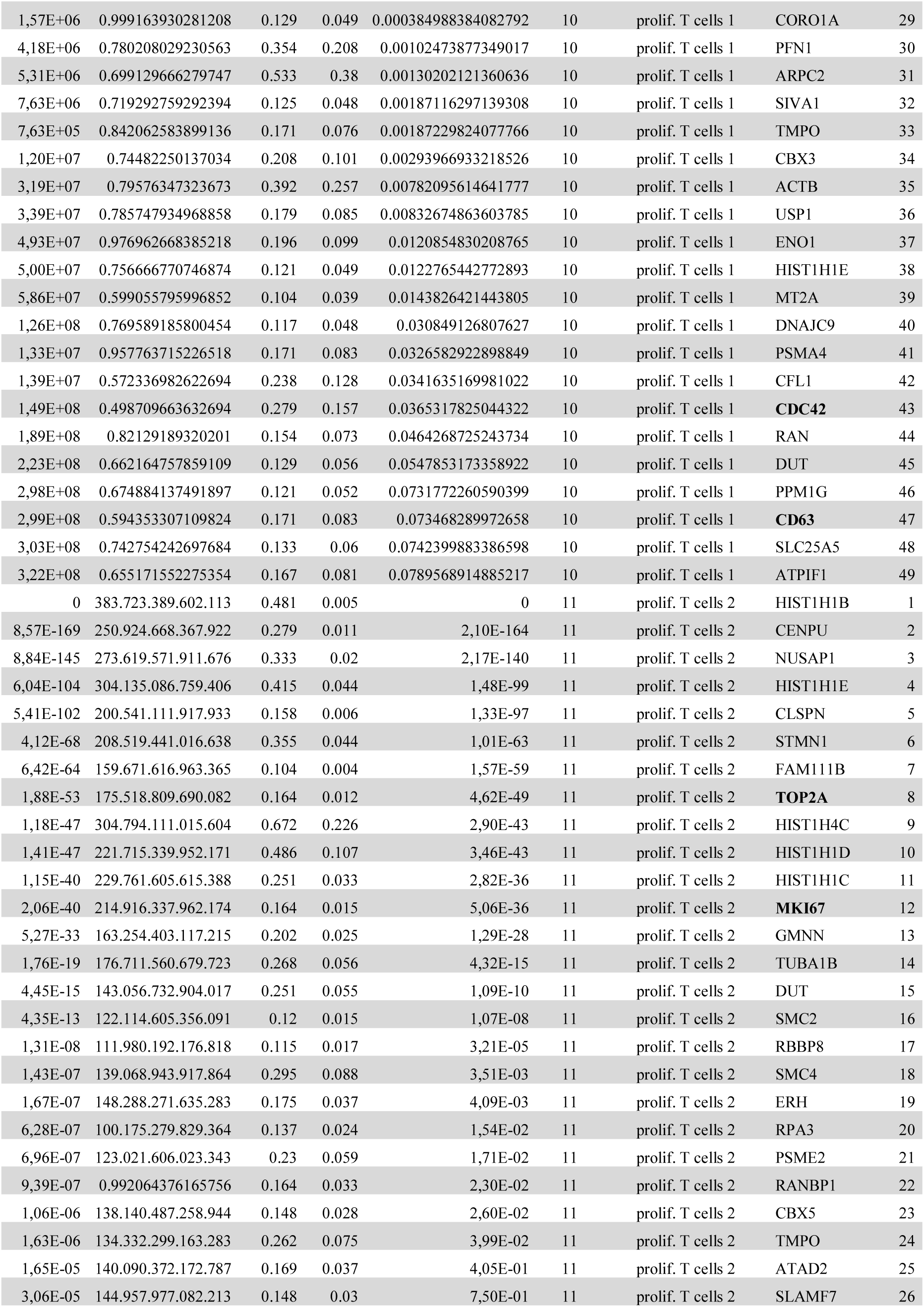
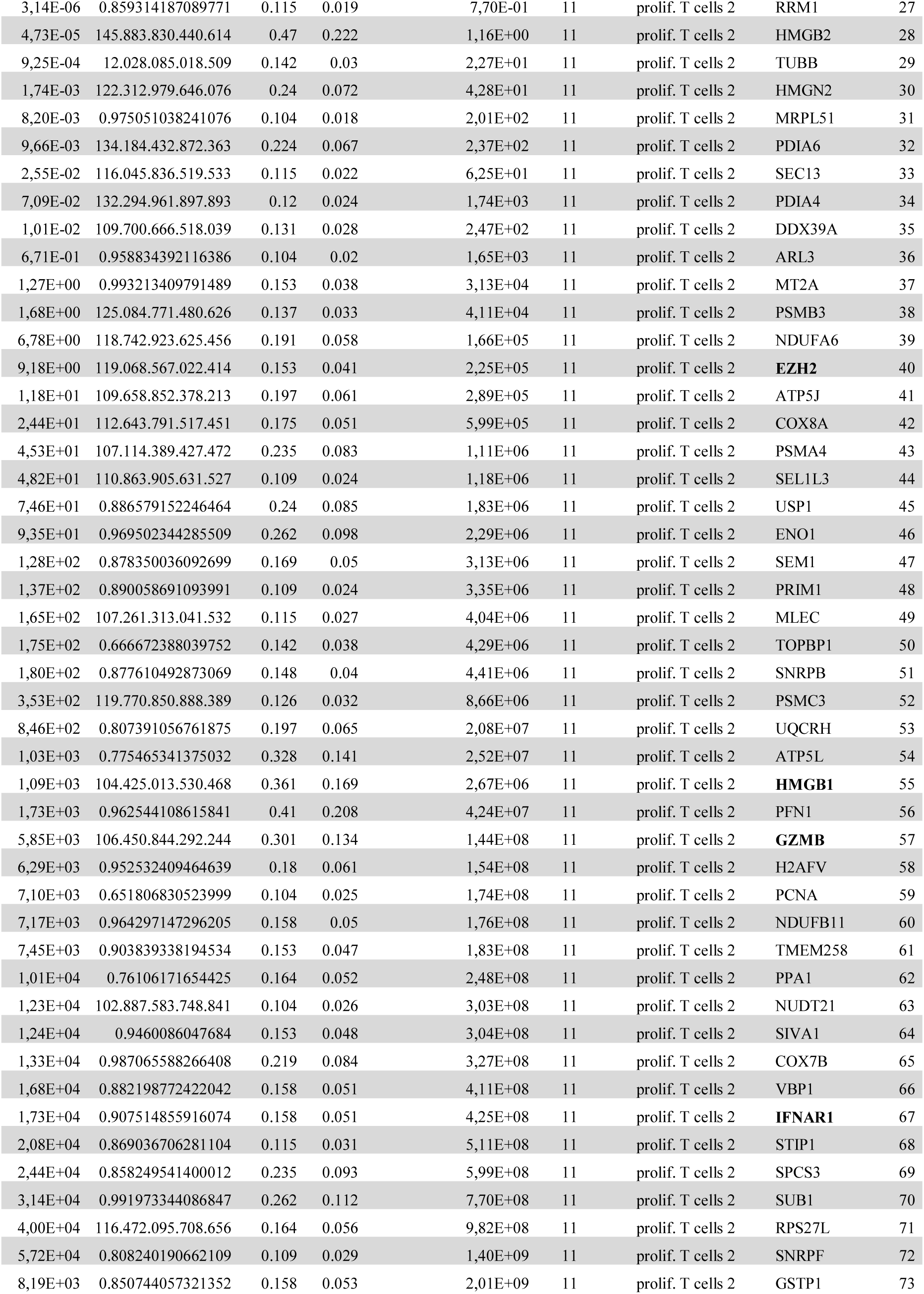
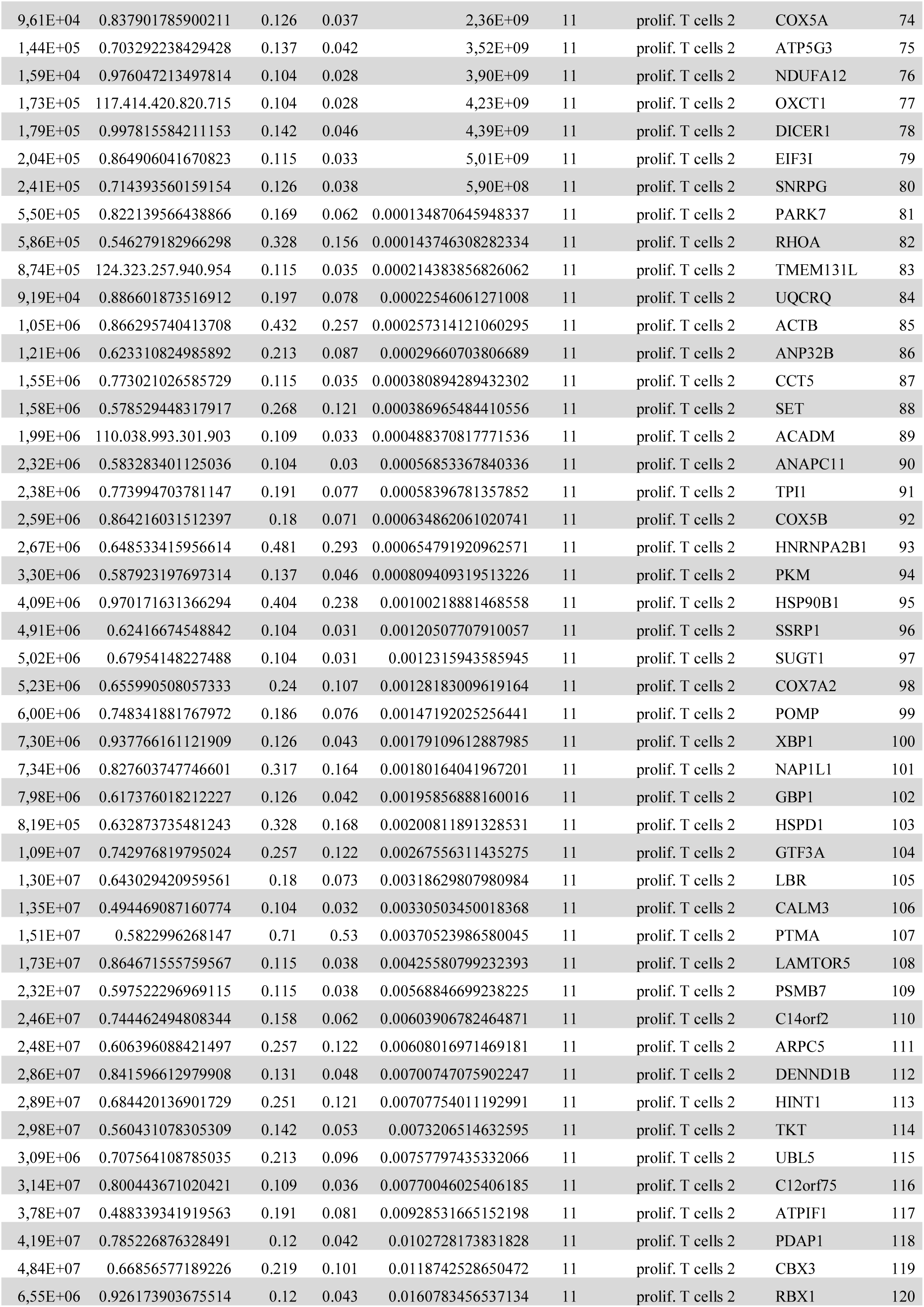
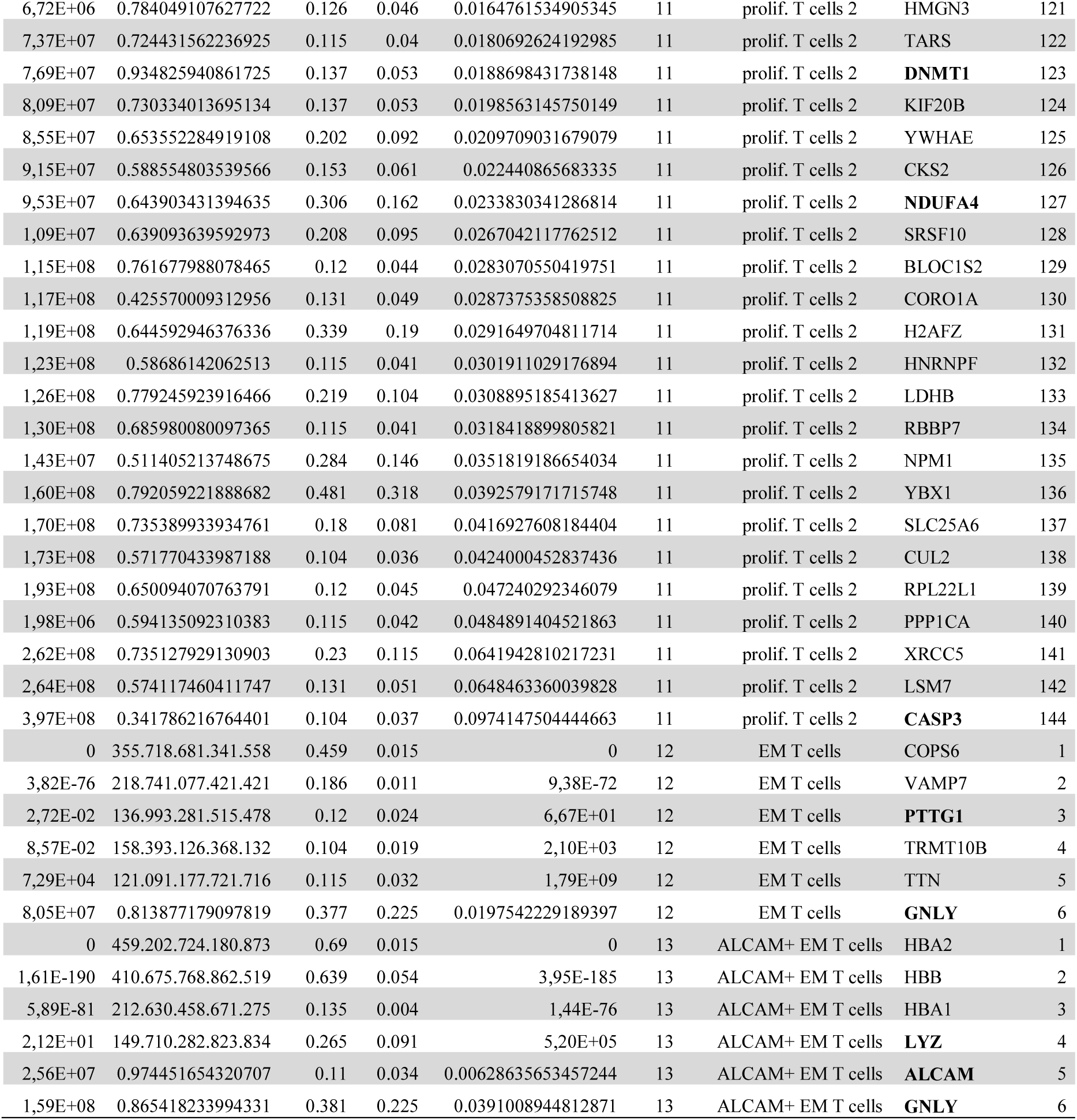
Differentially expressed genes.

