## Supplementary Material Bellmas-Sanz et al BioRxiv 2023 for "Donor tissue-resident memory-like T and NK cells generate a transient peripheral chimerism in lung transplant recipients, potentially protective from chronic lung allograft dysfunction"

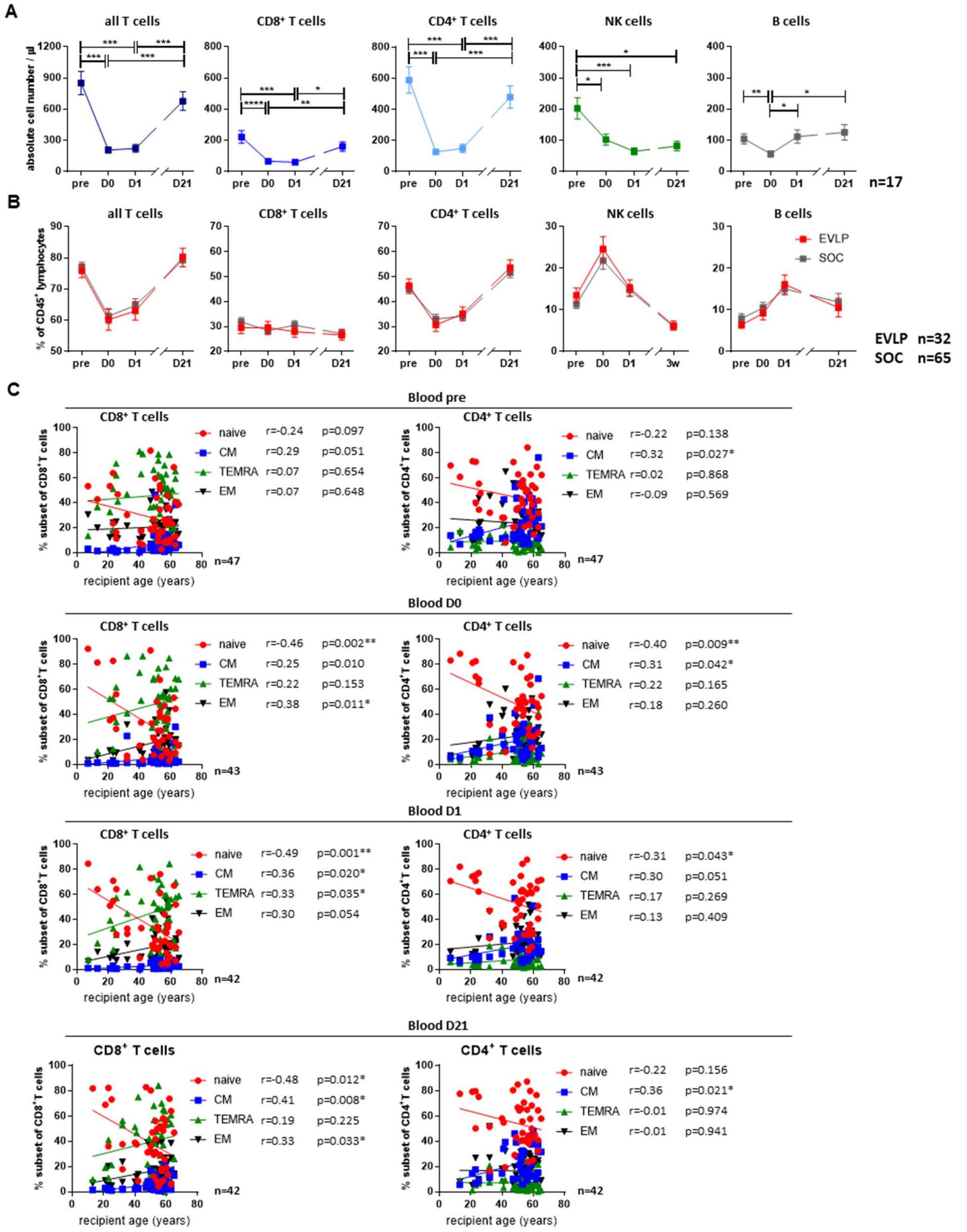


**Fig. S1. Absolute cell number all lymphocyte subsets decrease directly after transplantation. A**. Absolute cell number of all T cells, CD8$^+$ T cells, CD4$^+$ T cells, NK cells and B cells before transplantation (pre), and directly (D0), 24 hours (D1) and 3 weeks (D21) after transplantation (mean ± SEM, n=17). Cell numbers were calculated using BD Trucount Tubes. **B**. Proportions of lymphocyte subtypes at the indicated time points in patients that received a lung preserved under standard of care (SOC, n=65) or under Ex Situ Lung perfusion conditions (ESLP, n=32). Shown is mean ± SEM. **C**. Scatter plots showing linear regression between frequencies of $CD8^+$ and $CD4^+$ T cell subsets and recipient age before transplantation (pre, n=47) and at D0 (n=43), D1 (n=42) and D21 (n=42). Subsets were defined as naïve ($CD45RO^-CCR7^+$, red), CM ($CD45RO^+CCR7^+$, blue), TEMRA ($CD45RO^-CCR7^-$, green) and EM ($CD45RO^+CCR7^-$, black). Shown are the Pearson correlation coefficients (r) and their corresponding p-values (p). Statistical analysis: **A-B**. Repeated Measures ANOVA with Tukey multiple comparison test or Friedman test with Dunn's multiple comparison test. **C**. Pearson correlation. Asterisks indicate p values (*p<0.05, **p<0.01, ***p<0.001, ****p<0.0001).

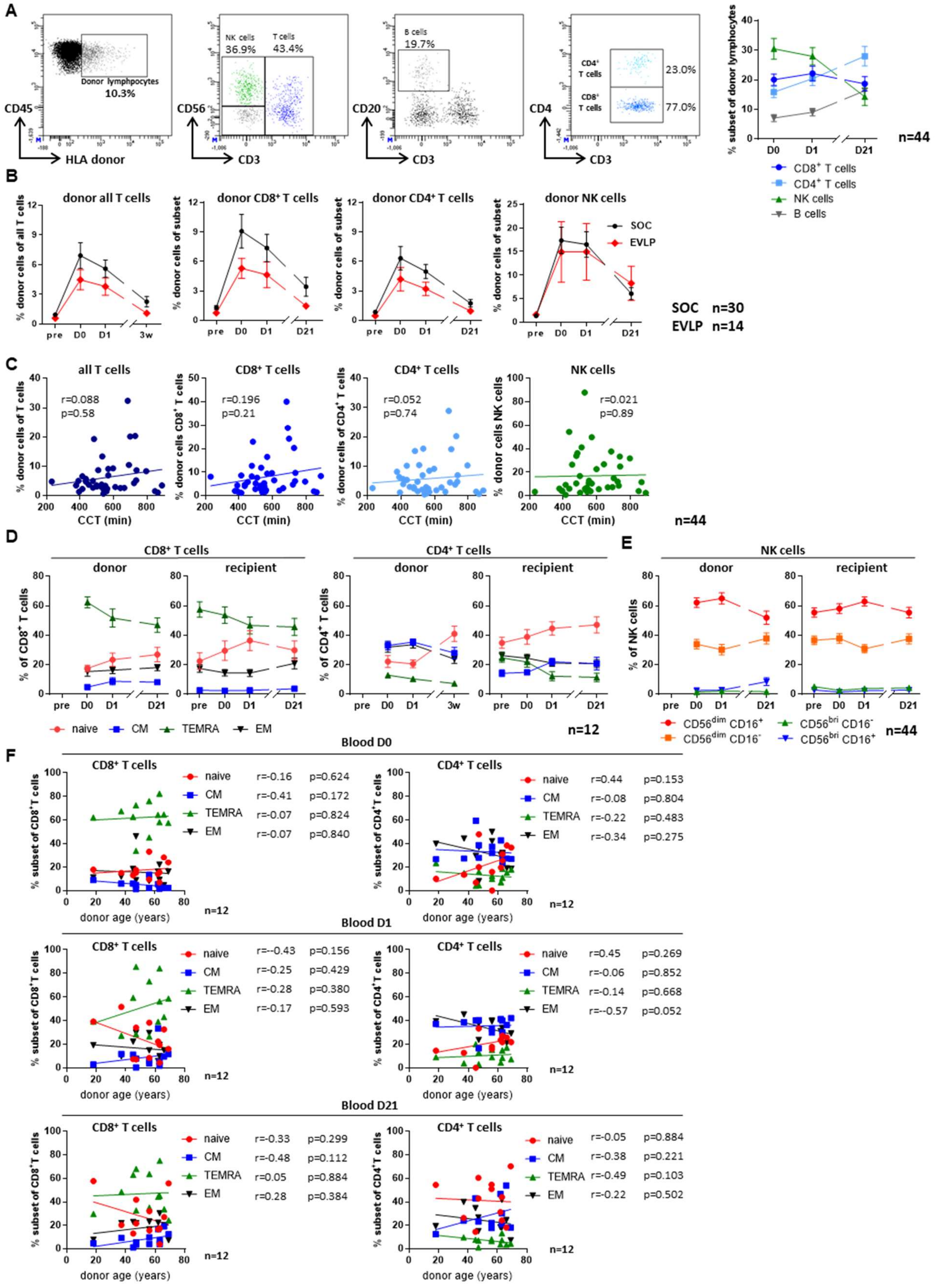


**Fig. S2**. **Further aspects on the dynamics of passenger donor lymphocyte subsets**. **A.** Representative flow cytometry plots and frequencies of donor CD8$^+$ T cells, CD4$^+$ T cells, NK cells and B cells among donor CD45$^+$ lymphocytes subsets before transplantation (pre), and directly (D0), 24 hours (D1) and 3 weeks (D21) after transplantation (mean ± SEM, n=44). **B**. Frequencies of donor cells among all T cells, CD8$^+$ T cells, CD4$^+$ T cells and NK cells at the indicated time points in patients that received a lung preserved under standard of care (SOC, n=30) or under Ex Situ Lung perfusion conditions (ESLP, n=12). Shown is mean ± SEM. **C**. Scatter plots showing linear regression between frequencies of donor T and NK and subsets and cross-clamp time (CCT) (n=44). Each dot represents one patient. Shown are the Pearson correlation coefficients (r) and their corresponding p-values (p). **D**. Proportions of CD8$^+$ and CD4$^+$ T cell subsets among donor and recipient T cells at the indicated time points (mean ± SEM, n=12). Subsets were defined as naïve (CD45RO$^-$CCR7$^+$, red), central memory (CD45RO$^+$CCR7$^+$, blue), TEMRA (CD45RO$^-$CCR7$^-$, green) and EM (CD45RO$^+$CCR7$^-$, black). **E**. Proportions of NK cells subsets among donor and recipient NK cells at the indicated time points (mean ± SEM, n=44). Subsets were defined as CD56$^{dim}$CD16$^+$ (red), CD56$^{dim}$CD16$^-$ (orange), CD56$^{bright}$CD16$^-$ (green) and CD56$^{bright}$CD16$^+$ cells (blue). **F**. Scatter plots showing linear regression between frequencies of donor CD8$^+$ and CD4$^+$ T cell subsets and donor age at D0, D1 and D21 (n=12). Shown are the Pearson correlation coefficients (r) and their corresponding p-values (p). Statistical analysis: **B**. Two-way repeated measures ANOVA with Sidak multiple comparisons test. **C, F**. Pearson correlation. Asterisks indicate p values (*p<0.05, **p<0.01, ***p<0.001, ****p<0.0001).

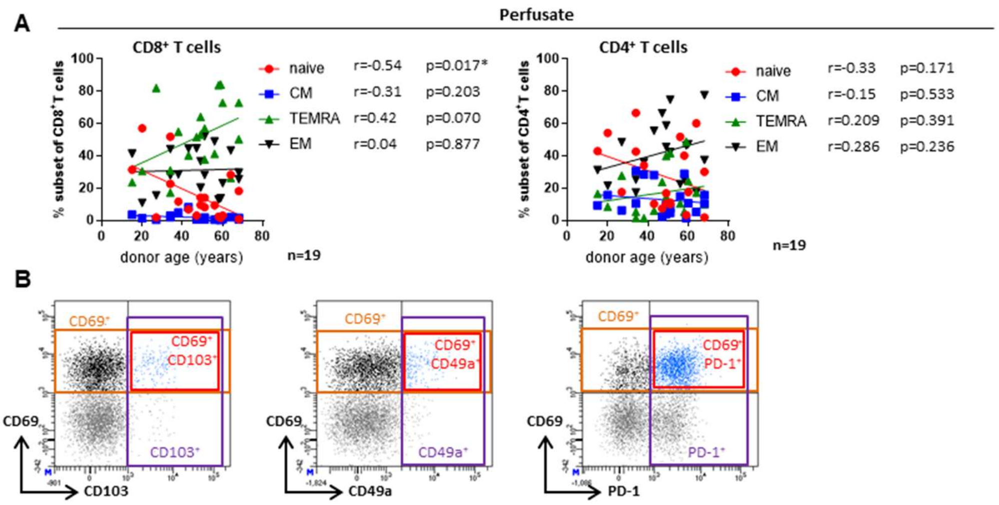


**Fig. S3. Correlation between donor age and T cell subsets in perfusates**. **A**. Scatter plots showing linear regression between frequencies of CD8$^+$ and CD4$^+$ T cell subsets in perfusate and donor age (n=19). Subsets were defined as naïve (CD45RO$^-$CCR7$^+$, red), central memory (CD45RO$^+$CCR7$^+$, blue), TEMRA (CD45RO$^-$CCR7$^-$, green) and EM (CD45RO$^+$CCR7$^-$, black). Shown are the Pearson correlation coefficients (r) and their corresponding p-values (p). Pearson correlation was performed, asterisks indicate p values (*p<0.05, **p<0.01, ***p<0.001, ****p<0.0001). **B**. Gating strategy for the identification of the different combinations of TRM markers (CD69, CD103, CD49a and PD-1).

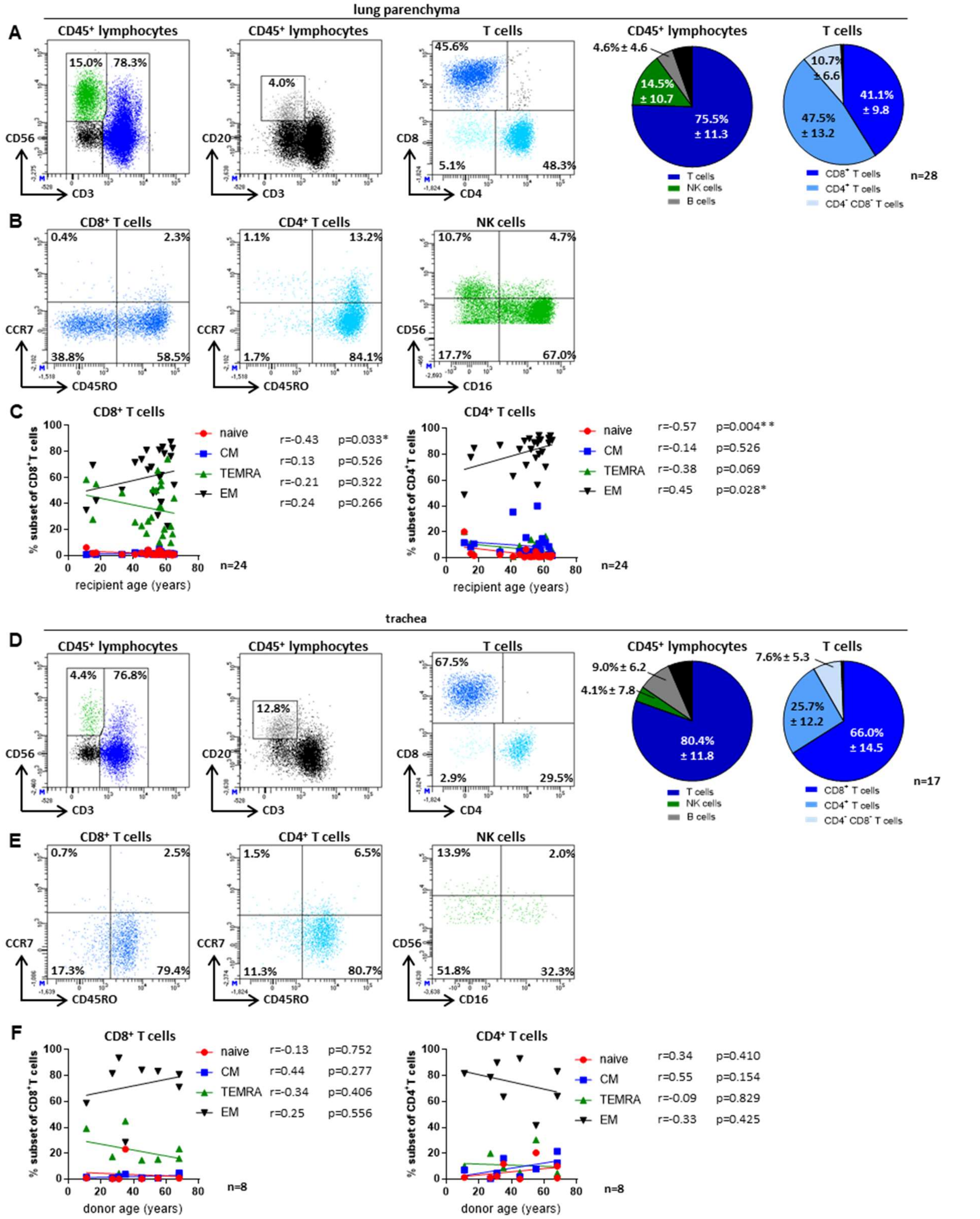


**Fig. S4. Lymphocyte composition of lung parenchyma and trachea. A**. Representative flow cytometry plots and frequencies of CD8$^+$ T cells, CD4$^+$ T cells, CD4$^-$CD8$^-$ T cells, NK cells and B cells in lung parenchyma (mean ± SD, n=28). **B**. Representative flow cytometry plots of CD8$^+$ T cell (n=24), CD4$^+$ T cell (n=24) and NK cell (n=26) subsets in lung parenchyma. T cells subsets include naïve (CD45RO-CCR7+), CM (CD45RO$^+$CCR7$^+$), TEMRA (CD45RO$^-$CCR7$^-$) and EM (CD45RO$^+$CCR7$^-$) populations. NK cell subsets include $CD56^{dim}CD16^{+}$, $CD56^{dim}CD16^{-}$, $CD56^{bright}CD16^{-}$ and $CD56^{bright}CD16^{+}$ populations. **C**. Scatter plots showing linear regression between frequencies of $CD8^{+}$ and $CD4^{+}$ T cell subsets in lung parenchyma and recipient age (n=24). Subsets were defined as naïve ($CD45RO^{-}CCR7^{+}$, red), central memory ($CD45RO^{+}CCR7^{+}$, blue), TEMRA ($CD45RO^{-}CCR7^{-}$, green) and EM ($CD45RO^{+}CCR7^{-}$, black). Shown are the Pearson correlation coefficients (r) and their corresponding p-values (p). **D**. Representative flow cytometry and frequencies of $CD8^{+}$ T cells, $CD4^{+}$ T cells, $CD4^{-}CD8^{-}$ T cells, NK cells and B cells in trachea (mean ± SD, n=17). **E**. Representative flow cytometry plots of $CD8^{+}$ T cell (n=24), $CD4^{+}$ T cell (n=24) and NK cell (n=26) subsets in the trachea. **F**. Scatter plots showing linear regression between frequencies of $CD8^{+}$ and $CD4^{+}$ T cell subsets in trachea and donor age (n=24). Shown are the Pearson correlation coefficients (r) and their corresponding p-values (p). Statistical analysis: **C-D**. Pearson correlation. Asterisks indicate p values (*p<0.05, **p<0.01, ***p<0.001, ****p<0.0001).

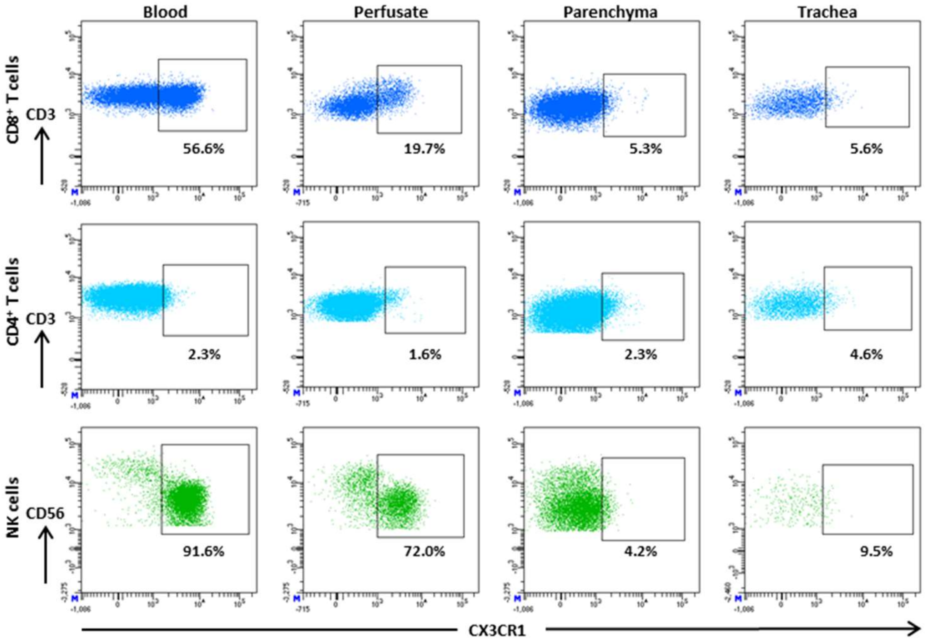


**Fig. S5**. **CX3CR1 is differentially expressed between the analyzed tissues a**. Representative flow cytometry plots showing CX3CR1 expression on $CD8^{+}$ T cells, $CD4^{+}$ T cells, NK cells and in blood before transplantation (pre), perfusate, lung parenchyma and trachea.
